# PRPS activity tunes redox homeostasis in Myc-driven lymphoma

**DOI:** 10.1101/2025.01.08.632009

**Authors:** Austin C. MacMillan, Bibek Karki, Juechen Yang, Karmela R. Gertz, Samantha Zumwalde, Jay G. Patel, Maria F. Czyzyk-Krzeska, Jarek Meller, John T. Cunningham

## Abstract

Myc hyperactivation coordinately regulates numerous metabolic processes to drive lymphomagenesis. Here, we elucidate the temporal and functional relationships between the medley of pathways, factors, and mechanisms that cooperate to control redox homeostasis in Myc-overexpressing B cell lymphomas. We find that Myc overexpression rapidly stimulates the oxidative pentose phosphate pathway (oxPPP), nucleotide synthesis, and mitochondrial respiration, which collectively steers cellular equilibrium to a more oxidative state. We identify Myc-dependent hyperactivation of the phosphoribosyl pyrophosphate synthetase (PRPS) enzyme as a primary regulator of redox status in lymphoma cells. Mechanistically, we show that genetic inactivation of the PRPS2 isozyme, but not PRPS1, in MYC-driven lymphoma cells leads to elevated NADPH levels and reductive stress-mediated death. Employing a pharmacological screen, we demonstrate how targeting PRPS1 or PRPS2 elicits opposing sensitivity or resistance, respectively, to chemotherapeutic agents affecting the thioredoxin and glutathione network, thus providing a therapeutic blueprint for treating MYC-driven lymphomas.

## Introduction

A master transcription factor capable of orchestrating both global and selective transcriptional responses^1–3^, the proto-oncogene *c-Myc* rewires metabolic processes en route to oncogenic transformation^4,5^. A well-established hallmark of Myc-dependent metabolic dysregulation is the altered utilization of nutrients such as glucose^6^. Glucose serves as a necessary precursor to fuel Myc-deregulated processes such as nucleotide biosynthesis^7^, RNA synthesis^8^, protein synthesis^9^ and biomass accumulation^10,11^. The global and selective upregulation of these processes is necessary to facilitate the increased anabolic demands of rapidly dividing Myc-overexpressing cells, as increased rates of cell cycle progression and maintenance of cell viability are intimately connected to the Myc-dysregulated metabolic program^12^. While individual Myc-deregulated pathways that contribute biosynthetic precursors, reducing equivalents and bioenergy are separately well-characterized and represent both dependencies and therapeutic vulnerabilities, a comprehensive understanding of their functional coordination in Myc-overexpressing malignancies remains elusive. As a prime example, an oxidative shift has been observed in different types of cancer^13^, as bioenergetic processes such as oxidative phosphorylation (OXPHOS) generate products that contribute to oxidative stress^14^. While this oxidative shift is appreciated as a consequence of Myc-driven tumorigenesis that can be leveraged to induce toxic levels of oxidative stress in tumor cells^15–17^, the primary mechanisms controlling the temporal and functional coordination between and among Myc-dependent processes that gives rise to altered redox homeostasis are still unclear.

In this study, we utilize multiple models of Myc-dependent lymphomagenesis to explore the mechanistic underpinnings of Myc-dysregulated redox homeostasis. In Myc-driven B cell lymphomas, overexpression is typically a consequence of a translocation event that repositions the *c-Myc* gene to the immunoglobulin heavy chain enhancer locus of chromosome 14^18^. We employ the Eμ-Myc mouse model^19^ and the P493-6 cell line^20^ to model the cellular response to supraphysiological Myc overexpression in B lymphocytes, we use splenic murine primary B lymphocytes activated with lipopolysaccharides (LPS) as a comparator to induce physiologically relevant enhanced levels of Myc expression, and we utilize the Burkitt’s lymphoma-derived CA46 and DG75 cell lines as models of bona fide Myc-dependent cancer cells. We compile a temporal atlas of the metabolic program under the direct control of oncogenic Myc and identify an early stimulation of oxidative metabolism as the functional consequence of an intricate connection between the mitochondrial electron transport chain (ETC), purine metabolism and the oxidative pentose phosphate pathway (oxPPP). We find that phosphoribosyl pyrophosphate synthetase (PRPS) enzymatic activity functionally links these redox processes, with evolutionarily conserved biochemical differences between the PRPS isoforms governing overall PRPS enzymatic efficiency. We discover that PRPS activity determines PPP flux, serving as an exit valve from the oxPPP and impacting global redox homeostasis in an isoform-specific manner. These metabolic changes are surprisingly uncoupled from Myc-regulated gene expression programs or anabolic processes, and we find that manipulating PRPP production via CRISPR/Cas9-based PRPS1 knockout (KO) or PRPS2 KO does not induce sensitivity to inhibitors of nucleotide metabolism. Rather, we demonstrate how inherent differences in PRPS1 and PRPS2 activity can be leveraged to elicit opposing effects on redox homeostasis to selectively target Myc-overexpressing lymphomas with compounds acting on key oxidizing or reducing machineries.

## Results

### Stimulation of oxidative metabolism is one of the earliest metabolic adaptations to Myc overexpression

We sought to determine the temporal organization of cellular responses (Fig. 1A) to Myc overexpression to better understand how Myc-dysregulated pathways are coordinated to promote global metabolic reprogramming. We first established a catalog of Myc-dysregulated molecular targets and pathways by performing RNA sequencing on murine primary B lymphocytes of the Eμ-Myc mouse model^19^, compared to wild-type (WT) mice (Fig. 1B). We next employed the P493-6 human cell line^20^ to temporally investigate the relationships between metabolic pathways that were identified as transcriptionally upregulated in the Eμ-Myc mice. We reasoned that a strategy of pairing functional consequences of Myc overexpression with the molecular contributors to the Myc overexpressing cellular phenotype could be used to more accurately define, order and group the key events in the Myc-dependent metabolic program. Consistent with Myc’s primary role as a transcription factor and in agreement with other studies^8^, one of the earliest events observed was an induction in global transcription within 2 hours after Myc induction, evident by increased total RNA content per cell (Supplementary Fig. 1A), levels of nucleotide biosynthetic enzymes and RNA Polymerase I, II and III levels and activation marks (Supplementary Fig. 1B). Interestingly, we identified a stark increase in mitochondrial respiration via oxygen consumption rate (OCR) (Fig. 1C) concurrent with increased nucleotide and RNA synthesis. While the OXPHOS-encoding genes have been identified as downstream targets of Myc in various contexts^21^, we observed that many individual components of the nuclear-encoded mitochondrial respiratory complexes maintained consistent levels of expression upon supraphysiological Myc induction (Supplementary Fig. 1C), suggesting that direct regulation of OXPHOS gene expression may not be the primary driver of oxidative metabolism. Additionally, we observed a gradual and sustained increase in both extracellular acidification rate (ECAR) (Fig. 1D) and glucose uptake (Fig. 1E, Supplementary Fig. 1D) over the time course of Myc overexpression which coincided with the Myc-dependent induction of the glycolytic enzymes hexokinase 2 (HK2) and lactate dehydrogenase (LDH) (Fig. 1E). This difference in the regulation of OXPHOS and glycolytic gene behavior is an indication that the Myc-dysregulated program is not an on/off switch, but rather a prioritized temporal activation of distinct interrelated metabolic processes.

**Figure 1.**
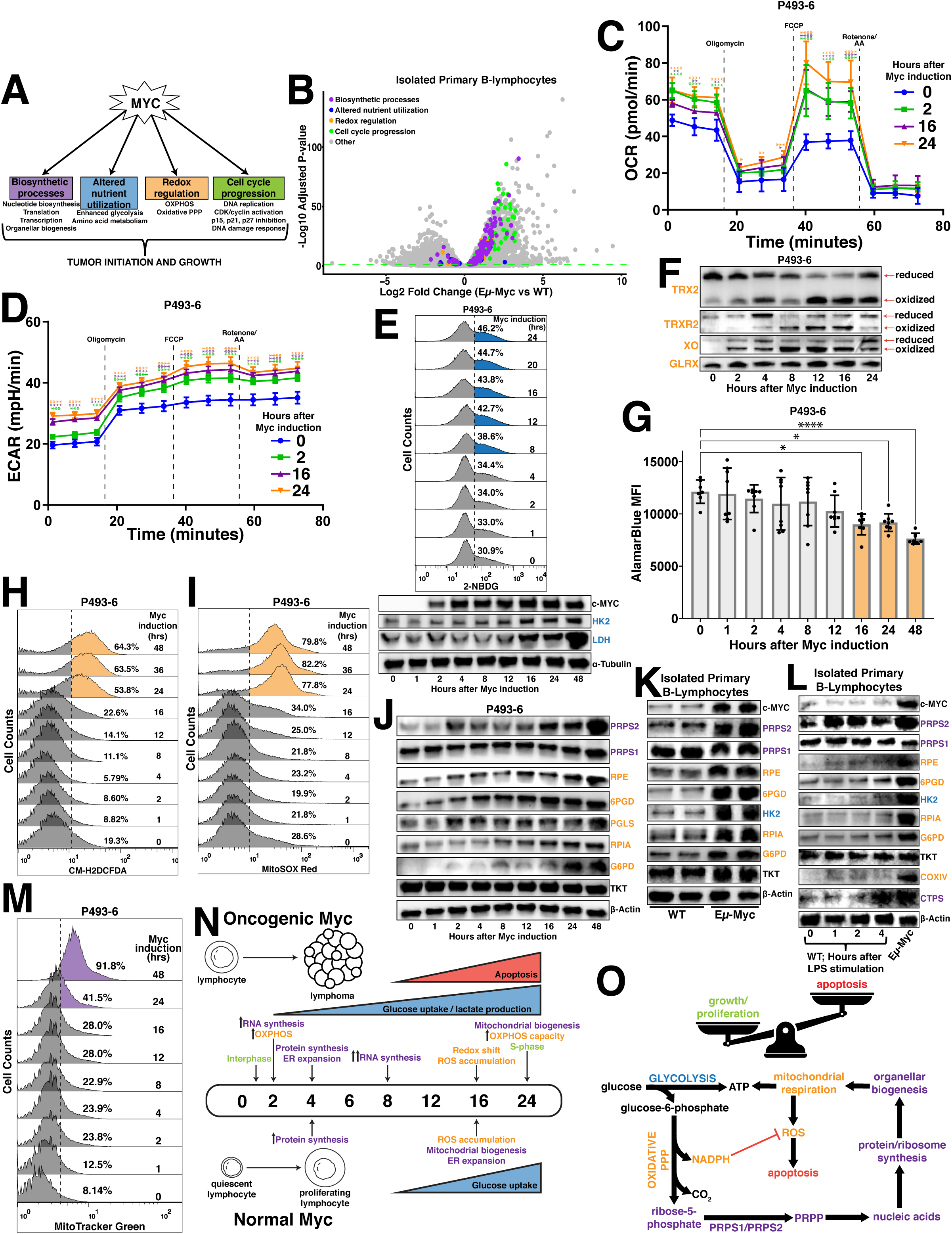
Myc overexpression stimulates oxidative metabolism. (A) Schematic depicting major Myc-dysregulated processes that contribute to tumor initiation and growth. (B) Volcano plot illustrating differentially expressed genes in Eµ-Myc vs wild-type (WT) murine primary B lymphocytes (Statistical analysis detailed in Methods; dashed green line demarks significance of p<0.05). (C) Oxygen consumption rate (OCR) and (D) extracellular acidification rate (ECAR) measured via Seahorse Mito Stress Test in P493-6 cells at times 0hr, 2hr, 16hr and 24hr following tetracycline removal to induce Myc expression. Dashed lines represent the time point at which oligomycin, FCCP and rotenone + antimycin A (AA) were added to the cells. (E) Glucose uptake in P493-6 cells over a 24hr time course following tetracycline removal to induce Myc expression, measured via 2-NBDG (top). Western blot of glycolytic enzyme expression in P493-6 cells over a 48hr time course following tetracycline removal to induce Myc expression. α-Tubulin used as a loading control (bottom). (F) Redox western blot illustrating protein oxidation state over a 48 hour time course following tetracycline removal to induce Myc expression. (G) AlamarBlue mean fluorescence intensity (MFI) as a readout of intracellular reduction in P493-6 cells over a 48hr time course following tetracycline removal to induce Myc expression. (H) Total intracellular reactive oxygen species (ROS) accumulation, measured via CM-H_2_DCFDA and (I) mitochondrial ROS accumulation, measured via MitoSOX Red in P493-6 cells over a 48hr time course following tetracycline removal to induce Myc expression. (J) Western blot of pentose phosphate pathway (PPP) metabolic enzyme expression in P493-6 cells over a 48hr time course following tetracycline removal to induce Myc expression. β-Actin used as a loading control. (K) Western blot of PPP metabolic enzyme expression in WT and Eµ-Myc murine primary B lymphocytes. β-Actin used as a loading control. (L) Western blot of PPP metabolic enzyme expression in WT murine primary B lymphocytes over a 4hr time course following LPS stimulation. Expression in Eµ-Myc murine primary B lymphocytes used for comparison. β-Actin used as a loading control. (M) Mitochondrial mass in P493-6 cells over a 48hr time course following tetracycline removal to induce Myc expression, measured via MitoTracker Green. (N) Timeline schematizing the temporal activation of metabolic processes following both oncogenic and normal levels of Myc induction. (O) Schematic depicting routes of glucose utilization to fuel metabolic processes that support lymphoma cell growth and proliferation downstream of oncogenic Myc. For all panels, statistical analysis performed via One-Way ANOVA. Bars represent mean ± s.d.; *p<0.05, **p<0.01, ***p<0.001, ****p<0.0001. For all histograms, upregulation quantified as a percentage of the population to the right of the dashed line at each time point.

4 hours after Myc induction, we observed significant increases in cytosolic translation rates (Supplementary Fig. 1E,F) that lagged the induced expression of well-established Myc targets eukaryotic translation initiation factor (eIF4E) and other ribosomal proteins (Supplementary Fig. 1G). Notably, induction of mitochondrial respiration and glycolytic flux preceded the increase in the protein synthesis rate, revealing distinct temporal patterns of Myc-dependent gene expression whereby at least some of the metabolic alterations downstream Myc are controlled by specific gene expression programs rather than being tethered to global increases in anabolism. Coincident with the induction of global translation rates, we observed an expansion of the endoplasmic reticulum (ER) compartment in the absence of an energy stress response (Supplementary Fig. 1H,I), which is consistent with previous findings that suggest a link between increased translation and activation of the unfolded protein response (UPR) in the ER^22^. Notably, murine primary B lymphocytes stimulated with LPS display increased rates of translation at a time point concurrent with that of the P493-6 cells (Supplementary Fig. 1J), but required more time to induce ER expansion and mitochondrial biogenesis (Supplementary Fig. 1K,L). These data establish transcription and stimulation of oxidative metabolism to be the earliest adaptations to Myc-dysregulated metabolism, closely followed by enhanced global translation and organellar biogenesis that occurs after an earlier specific translational response.

### Myc-dependent oxidative metabolism overwhelms the reductive potential of oxPPP-generated NADPH

The early Myc-dependent upregulation of OXPHOS led us to hypothesize that increased respiration may alter intracellular redox homeostasis. To test this, we first performed redox western blotting to assay the oxidative state of xanthine oxidase (XO), a component of the bifunctional xanthine oxidoreductase (XOR) enzyme which oxidizes hypoxanthine during purine catabolism, as well as the mitochondrial-localized thioredoxin 2 (TRX2) and thioredoxin reductase 2 (TRXR2), which utilize the reducing equivalent NADPH to alleviate respiration-linked oxidative stress. We observed a distinct oxidation of reactive cysteines in each these proteins at a time point concurrent with the induction of glucose uptake, though the lack of such shift in cysteine oxidation of the glutathione-dependent glutaredoxin (GLRX) protein hints at selectivity within the Myc-regulated oxidative program (Fig. 1F). We next utilized the AlamarBlue dye as a bio-orthogonal measure of intracellular reductive activity, as the conversion of resazurin to the fluorescent molecule resorufin is primarily reliant upon NAD(P)H-dependent oxidase activity. We did not observe a statistically significant decrease in AlamarBlue fluorescence until 16 hours post-Myc induction (Fig. 1G), which was concomitant with an increase in total intracellular and mitochondrial reactive oxygen species (ROS) in both the P493-6 cells (Fig. 1H,I) and LPS-stimulated WT murine primary B lymphocytes (Supplementary Fig. 1M). Despite the pro-apoptotic factor Bim being among the earliest Myc targets to be induced, observable increases in apoptosis via poly[ADP-ribose] polymerase 1 (PARP1) and caspase 3 cleavage are not evident until 12-16 hours after supraphysiological Myc overexpression (Supplementary Fig. 1Q), which coincides with the appearance of indicators of oxidative stress and the shift in redox homeostasis. Interestingly, our data illustrates that the Myc-dependent decrease in AlamarBlue reduction is concurrent with the upregulation of kelch-like ECH-associated protein 1 (KEAP1) and downregulation of nuclear factor erythroid 2-related factor 2 (NRF2) targets heme oxygenase-1 (HO-1), NAD(P)H:quinone oxidoreductase 1 (NQO1), thioredoxin reductase 1 (TRXR1), TRXR2 and glutathione reductase (GSR) while other established NRF2 targets such as superoxide dismutase 2 (SOD2), thioredoxin 1 (TRX1), catalase, glutamate-cysteine ligase modifier subunit (GCLM) and peroxiredoxin 1 (PRDX1) display no change in expression over the time course of Myc overexpression (Supplementary Fig. 1N), suggesting the NRF2 pathway is not a primary determinant of redox homeostasis under these conditions^23^. Additionally, our data indicates that dehydrogenase-mediated reactions of the tricarboxylic acid (TCA) cycle and folate metabolism are not major contributors to Myc-driven oxidative metabolism, as a majority of the enzymes involved in these pathways display no pattern of upregulation in response to supraphysiological Myc overexpression (Supplementary Fig. 1O). Because the oxPPP links glucose metabolism and nucleotide biosynthesis while serving as a major source of NADPH production^24^ and has been shown to be Myc-regulated in other cancers^5,6^, we next sought to assess its potential dysregulation downstream of Myc overexpression. Indeed, we observed an early induction of the PPP enzymes ribulose-5-phosphate-3-epimerase (RPE), 6-phosphogluconate dehydrogenase (6PGD) and 6-phosphogluconolactonase (PGLS) with later induced expression of ribose 5-phosphate isomerase A (RPIA) and glucose-6-phosphate dehydrogenase (G6PD), while the expression of the non-oxPPP enzyme transketolase (TKT) displayed no such upregulation (Fig. 1J). While these findings are consistent in pre-malignant Eμ-Myc murine primary B lymphocytes (Fig. 1K), they are dependent upon supraphysiological levels of Myc expression, as LPS-stimulated WT primary B lymphocytes do not display the same early induction or magnitude of oxPPP gene expression (Fig. 1L). Moreover, the Myc-dependent increase of the NADPH-generating oxPPP enzymes G6PD and 6PGD occurred in the absence of a corresponding increase in transcripts encoding those oxPPP enzymes, implicating transcription-independent roles for Myc during metabolic reprogramming. Importantly, the metabolic effects governing this oxidative shift precede later Myc-dependent changes to mitochondrial biogenesis, as we did not observe a significant accumulation of mitochondrial mass (Fig. 1M) until 24 hours after Myc induction, which coincided with an increase in maximum respiratory capacity (Fig. 1C) and S-phase progression (Supplementary Fig. 1P,Q). However, not all organellar biogenesis processes are upregulated downstream of Myc, as there was a slight diminishment of lysosomal content following supraphysiological Myc induction (Supplementary Fig. 1R). Collectively, this analysis establishes a timeline of Myc-dependent metabolic reprogramming (Fig. 1N), where near-immediate activation of the oxPPP directs glucose to nucleotide biosynthesis to facilitate early anabolic processes and provides the reducing equivalent NADPH that can be used to both counteract ROS and augment ETC activity via reactions catalyzed by enzymes such as inosine monophosphate dehydrogenase (IMPDH) and XOR. These results fit a model whereby these metabolic processes sustain lymphoma cell growth and proliferation, which outcompetes the apoptotic phenotype that arises when the reducing capacity generated via oxPPP-mediated NADPH production becomes overwhelmed by the oxidative byproducts of other pathways engaged upon supraphysiological Myc overexpression (Fig. 1O).

### Mitochondrial respiration, purine cycling and oxPPP are intrinsically linked to promote Myc-driven oxidative metabolism

To elucidate which processes are functionally responsible for the increased oxidative metabolism downstream of Myc, we treated P493-6 cells with inhibitors of pathways previously linked to redox homeostasis (Fig. 2A) and conducted a temporal analysis of AlamarBlue reduction. We inhibited OXPHOS in the P493-6 cells by treating with chloramphenicol because the translation of key redox-regulating components of the ETC are mitochondrially-encoded^25,26^; we inhibited oxPPP activity by treating with G6PDi-1^27^ because the oxPPP is a major source of NADPH production and a link has been established between Myc and the oxPPP in some cancers^28^; we inhibited XOR by treating with allopurinol because of its redox bifunctionality as an oxidoreductive enzyme^29^ and the studies that have uncovered a connection between purine regulation and mitochondrial metabolism^30,31^; and, we inhibited dihydroorotate dehydrogenase (DHODH) with brequinar because of its function as a mitochondrial membrane-localized oxidoreductive enzyme that transfers electrons to ubiquinol during *de novo* pyrimidine biosynthesis. We observed that treatment with chloramphenicol, G6PDi-1 and allopurinol completely abrogated the Myc-dependent decrease in AlamarBlue fluorescence upon supraphysiological Myc overexpression, whereas treatment with brequinar failed to do so (Fig. 2B). Together, these results pinpoint OXPHOS, purine metabolism and the oxPPP as the major functional determinants of a Myc-driven oxidative program, as inhibiting any of these processes abrogates the oxidative shift observed upon Myc induction.

**Figure 2.**
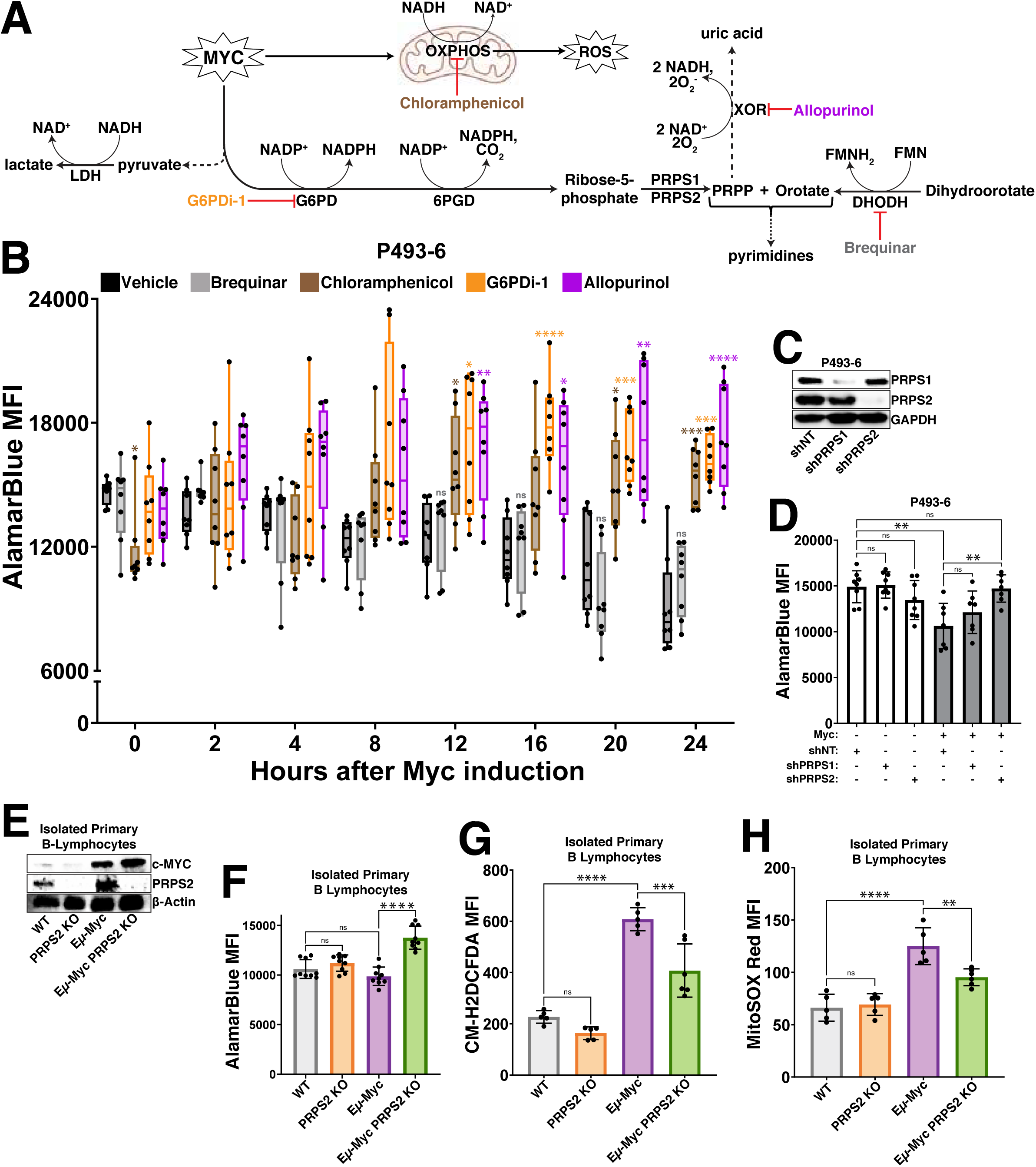
Myc-dependent coordination of OXPHOS, oxPPP and purine metabolism dictates redox state. (A) Schema for inhibiting major regulators of redox homeostasis downstream of Myc overexpression. (B) AlamarBlue mean fluorescence intensity (MFI) as a readout of intracellular reduction in P493-6 cells treated with vehicle control (DMSO), brequinar, chloramphenicol, G6PDi-1 or allopurinol over a 24hr time course following tetracycline removal to induce Myc expression. (C) Western blot validation of shRNA-mediated knockdown of PRPS1 and PRPS2 in P493-6 cells. GAPDH used as a loading control. (D) AlamarBlue MFI as a readout of intracellular reduction in P493-6 cells under different conditions (Myc OFF, Myc OFF shPRPS1, Myc OFF shPRPS2; Myc ON, Myc ON shPRPS1, Myc ON shPRPS2). (E) Western blot validation of murine primary B lymphocyte genotypes. β-Actin used as a loading control. (F) AlamarBlue MFI as a readout of intracellular reduction in murine primary B lymphocytes of the indicated genotypes. (G) Total intracellular ROS accumulation, measured via CM-H_2_DCFDA MFI and (H) mitochondrial ROS accumulation, measured via MitoSOX Red MFI in murine primary B lymphocytes of the indicated genotypes. For all panels, statistical analysis performed via One-Way ANOVA, bars represent mean ± s.d.; *p<0.05, **p<0.01, ***p<0.001, ****p<0.0001, ns: not significant.

We next sought to determine the metabolic node connecting these redox-regulating processes, where we nominated the PRPS isozymes (PRPS1, PRPS2), which catalyze the conversion of the oxPPP product ribose-5-phosphate (R5P) to the nucleotide biosynthesis pathway intermediate phosphoribosyl pyrophosphate (PRPP)^32^. We chose these enzymes because of the early induction of PRPS2 expression in response to both supraphysiological and normal levels of Myc overexpression (Fig. 1J-L) and the crucial role of PRPS2 in maintaining lymphoma cell viability^33^. We first confirmed that the PRPS isozymes co-assemble, together with the PRPS associated proteins (PRPSAP1, PRPSAP2), to form a complex^34^ in Myc-dependent lymphoma cells (Supplementary Fig. 2A). Interestingly, we observed that the Myc-dependent induction of PRPS2 expression altered PRPS complex configuration, resulting in the downshift of a large molecular weight complex and an enrichment of a smaller molecular weight dimeric configuration between PRPS1 and PRPS2 (Supplementary Fig. 2B). Importantly, this dimeric PRPS configuration suggests a loss of purine-mediated allosteric feedback inhibition, as three subunits of PRPS isozymes are required to create an allosteric binding pocket^35^. To test whether isozyme-dependent remodeling of the PRPS enzyme complex is functionally important for Myc-dependent oxidative metabolism, we performed shRNA-mediated knockdown of PRPS1 and PRPS2 expression in P493-6 cells (Fig. 2C) and compared AlamarBlue reduction +/- Myc. We observed no significant change to AlamarBlue conversion upon knockdown of either PRPS isoform in cells expressing low levels of Myc, but the decreased levels of AlamarBlue reduction upon supraphysiological Myc overexpression were completely rescued upon knockdown of PRPS2, but not PRPS1 (Fig. 2D). These results were recapitulated in murine primary B lymphocytes (Fig. 2E), where we saw a similar decrease of AlamarBlue reduction (Fig. 2F) paired with significantly elevated levels of both total intracellular (Fig. 2G) and mitochondrial (Fig. 2H) ROS in Eμ-Myc B cells compared to Eμ-Myc; PRPS2 KO B cells. However, no significant change was observed between WT and PRPS2 KO B cells without Myc overexpression. Our previous work found there to be no significant differences between either mitochondrial mass or membrane potential upon PRPS2 KO in either WT or Eμ-Myc B lymphocytes^33^, suggesting that the Myc-dependent oxidative program does not require mitochondrial biogenesis but rather activity of the PRPS complex. Together, these data identify OXPHOS, purine metabolism and oxPPP activity as the critical circuitry controlling redox homeostasis in Myc-overexpressing lymphomas and nominate PRPS activity as a central hub coupling these cytosolic and mitochondrial redox processes.

### PRPS activity couples viability and mitochondrial respiration in Myc-driven lymphoma

To assess the requirement of Myc-driven, PRPS-dependent redox regulation in the context of fully transformed B cell lymphoma, we generated single-cell selected CRISPR/Cas9 knockouts of both PRPS1 and PRPS2 in CA46 and DG75 Burkitt’s lymphoma-derived cell lines (Fig. 3A). The drastic increase in the cleavage of PARP1 in the PRPS2 KO cells indicates they are significantly more apoptotic, aligning with our previous findings^33^. This phenotype is specific to the Myc-regulated PRPS2 isoform, as knocking out the PRPS1 isoform does not impact viability in these lymphoma cells, suggesting that the Myc-dependent induction of oxidative metabolism may be linked to lymphoma cell viability via PRPS activity. To define the redox alterations upon PRPS2 KO, we measured the levels of key indicators of redox state including NADPH/NADP+ (Supplementary Fig. 2C,D), NADH/NAD+ (Supplementary Fig. 2E-G) and GSH/GSSG (Supplementary Fig. 2H-J). We observed a significant increase in NADPH levels (Fig. 3B), reduced glutathione levels (GSH) (Fig. 3C) and AlamarBlue reduction (Fig. 3D) in the PRPS2 KO cells of both cell lines, while the levels of both total intracellular and mitochondrial ROS were significantly decreased (Fig. 3E,F; Supplementary Fig. 3A,B). We completely restored both the AlamarBlue reduction and viability in the PRPS2 KO cells of each cell line upon exogenous overexpression of NDI1, a single polypeptide encoding the yeast NADH-quinone oxidoreductase enzyme to augment ETC Complex I activity^36^, as well as alternative oxidase (AOX), which encodes the terminal oxidase for the ETC in plants^37^, indicating a functional coupling between cytosolic and mitochondrial redox networks. We were also able to rescue the NADPH-induced reductive stress of the PRPS2 KO cells with exogenous expression of cytosolic or mitochondrial triphosphopyridine nucleotide oxidase (TPNOX), an engineered *Lactobacillus brevis*-derived oxidase designed to consume intracellular NADPH by catalyzing the reaction 2NADPH + 2H^+^ + O_2_ → 2NADP^+^ + 2H_2_O^38^ (Fig. 3G,H). Together, these data confirm an oxidative program in fully transformed Myc-driven lymphoma and suggest that viability and oxidative metabolism are intrinsically linked via PRPS activity.

**Figure 3.**
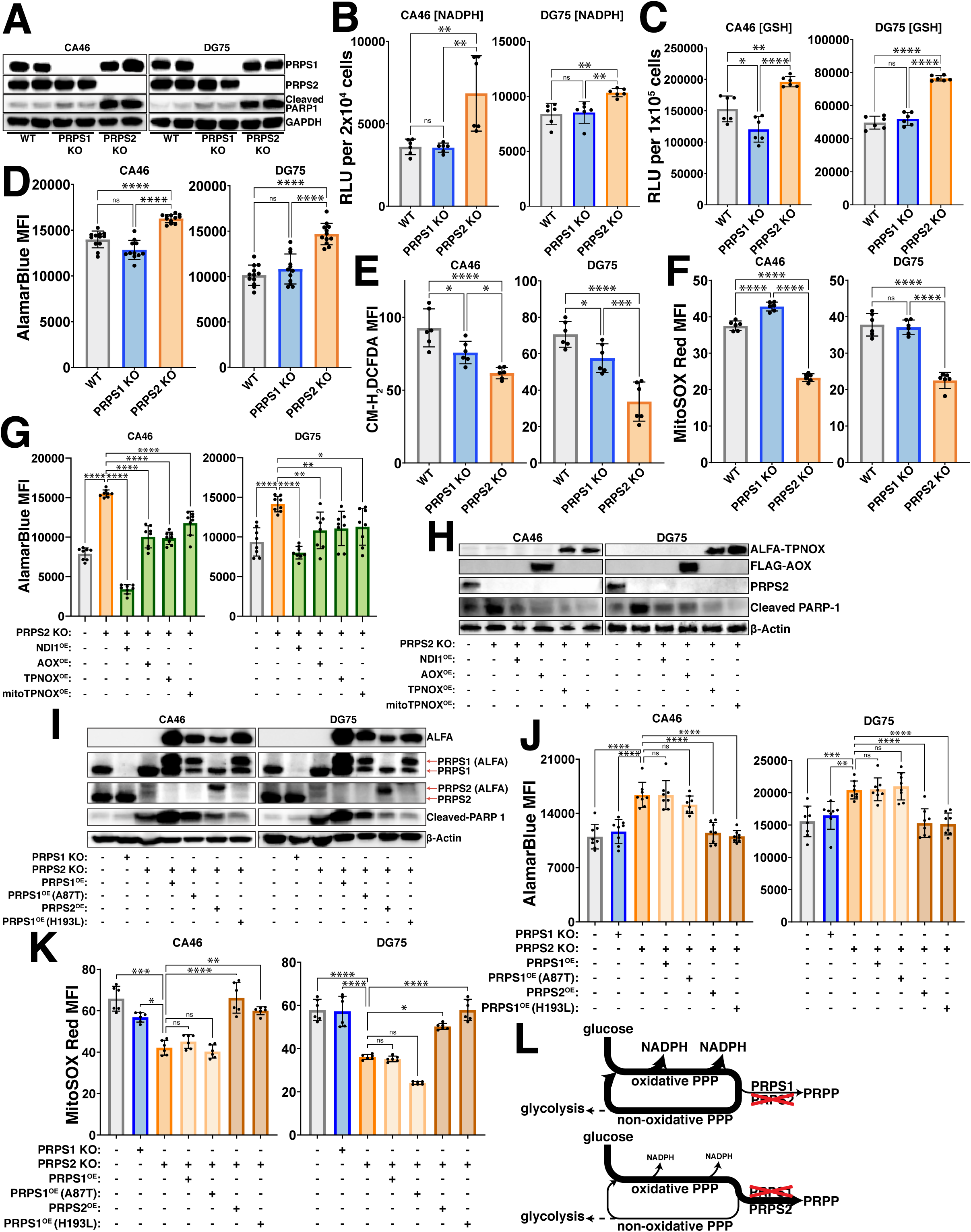
PRPS activity governs redox balance. (A) Western blot validating CRISPR/Cas9 mediated knockout of PRPS1 and PRPS2 in two separate human Burkitt’s lymphoma-derived cell lines, CA46 (left) and DG75 (right). 24 kDa PARP-1 fragment is used as an apoptotic marker. GAPDH used as a loading control. Levels of (B) NADPH and (C) reduced glutathione (GSH) in WT, PRPS1- and PRPS2-KO cells of CA46 (left) and DG75 (right) cell lines, measured via relative luciferase units (RLU) of luminescent-based GLO-assays. (D) AlamarBlue mean fluorescence intensity (MFI) as a readout of intracellular reduction in WT, PRPS1- and PRPS2- KO cells of CA46 (left) and DG75 (right) cell lines. (E) Total intracellular ROS accumulation, measured via CM-H_2_DCFDA MFI and (F) mitochondrial ROS accumulation, measured via MitoSOX Red MFI in WT, PRPS1- and PRPS2- KO cells of CA46 (left) and DG75 (right) cell lines. (G) AlamarBlue MFI as a readout of intracellular reduction and in WT, PRPS2 KO and PRPS2 KO cells containing exogenously expressed NDI1, AOX, TPNOX or mitoTPNOX of CA46 (left) and DG75 (right) cell lines. (H) Western blot of PARP-1 cleavage as an apoptotic marker in WT, PRPS2 KO and PRPS2 KO cells containing exogenously expressed NDI1, AOX, TPNOX or mitoTPNOX of CA46 (left) and DG75 (right) cell lines. β-Actin used as a loading control. (I) Western blot validating stable exogenous expression of ALFA-tagged PRPS1, PRPS2 and PRPS1 hypomorphic (A87T) and superactive (H193L) mutant constructs in CA46 (left) and DG75 (right) PRPS2 KO cell lines. 24 kDa PARP-1 fragment is used as an apoptotic marker. β-Actin used as a loading control. (J) AlamarBlue MFI as a readout of intracellular reduction and (K) MitoSOX Red MFI as a readout of mitochondrial ROS accumulation in WT, PRPS1 KO, PRPS2 KO and PRPS2 KO cells containing stably integrated ALFA-tagged PRPS1, PRPS2, PRPS1 hypomorphic mutant (A87T) and PRPS1 superactive mutant (H193L) constructs in CA46 (left) and DG75 (right) cell lines. (L) Schematic depicting the escape or trapping of glucose-derived carbons in a PPP cycle, governed via PRPS enzymatic efficiency. For all panels, statistical analysis performed via One-Way ANOVA, bars represent mean ± s.d.; *p<0.05, **p<0.01, ***p<0.001, ****p<0.0001, ns: not significant.

### Feedback-refractory biochemical property of PRPS2 promotes the Myc-driven oxidative program

Though the PRPS isoforms are evolutionarily conserved and exhibit 95% amino acid homology in humans^34^, *in vitro* studies have shown that recombinant PRPS1 isoform is much more sensitive than PRPS2 to inhibition by downstream purine products^32^, an allosteric feedback regulatory mechanism found in Class I PRPS enzymes conserved from their bacterial origins^39^. Several disease-causing superactivating mutations in the X-linked *PRPS1* gene render the PRPS enzyme feedback-refractory to purine-mediated allosteric inhibition have been identified in humans^40^. Of note, the heterozygous H193L mutation was identified in a young female patient, confirming its pathogenicity in the presence of WT PRPS1 and PRPS2 alleles^41^. To test whether differences in allosteric feedback sensitivity to purines account for the phenotypes observed in PRPS2 KO lymphoma cells, we generated ALFA-tagged PRPS1 harboring superactive D52H and H193L mutations to exogenously overexpress in PRPS2 KO cells. To test the alternative hypothesis that PRPS2 upregulation controls enzymatic efficiency of the PRPS complex through structural alterations that are independent of its own enzymatic activity, we engineered a catalytically inactive PRPS2 E39A mutation which renders the enzyme non-functional by abrogating hydrogen bonding between the adenine N6 atom of ATP and the side chain of the evolutionarily conserved glutamic acid at the active site^34^. As an additional negative control, we employed an ALFA-tagged hypomorphic PRPS1 A87T variant which has decreased ATP binding affinity and causes congenital sensorineural hearing loss (DFN2) in humans^42^. Exogenous expression of ALFA-PRPS2 or either of the superactive ALFA-PRPS1 D52H/H193L mutants was sufficient to rescue the viability (Fig. 3I, Supplementary Fig. 3C) and restore the levels of AlamarBlue reduction (Fig. 3J, Supplementary Fig. 3D) and mitochondrial ROS accumulation (Fig. 3K; Supplementary Fig. 3E,F) of PRPS2 KO lymphoma cells, whereas exogenous expression of ALFA-PRPS1, hypomorphic ALFA-PRPS1 A87T or catalytically inactive ALFA-PRPS2 E39A did not. These data support a model by which supraphysiological Myc overexpression selectively upregulates PRPS2, resulting in PRPS2-dependent remodeling of the PRPS complex which feeds oxPPP-derived carbons into purine production and catabolism pathways, to promote OXPHOS and establish and maintain an oxidative metabolic program (Fig. 3L).

### PRPS-dependent control of redox state is uncoupled from Myc-dependent anabolic processes or cell cycle progression

Our temporal analyses indicate that Myc’s early gene expression program promotes increased oxPPP flux and PRPP-dependent nucleotide production coincident with an increase in RNA synthesis, so we questioned whether and how PRPS activity is linked to Myc-regulated control of transcription and other anabolic processes. We first performed RNA-sequencing in the primary B lymphocytes from pre-malignant Eµ-Myc and Eµ-Myc; PRPS2 KO mice. Of the 4 sufficiently expressed genes found to be differentially regulated (log_2_(fold change)≥0.95), *Prps2* is the only gene that notably exceeded the threshold for significance (p<0.05) (Fig. 4A). To test whether PRPS2 KO could counteract Myc’s role as global regulator of anabolic metabolism in bona fide models of fully transformed Myc-driven lymphoma, we measured total RNA content per cell (Fig. 4B), protein synthesis rates (Supplementary Fig. 4A), total protein per cell (Fig. 4C), average cell diameter (Fig. 4D) and mitochondrial mass (Fig. 4E), and observed no significant changes upon PRPS1 KO or PRPS2 KO in CA46 and DG75 cells. ER-tracker staining revealed a decrease in ER content in both PRPS1 KO and PRPS2 KO cells compared to WT, with no significant changes between either KO cell line (Fig. 4F). We assayed the cell cycle phase occupancy of both lymphoma cell lines and the murine primary B lymphocytes and observed no consistent changes between cell cycle profiles upon PRPS1 KO or PRPS2 KO (Fig. 4G, Supplementary Fig. 4B), demonstrating that the loss of either PRPS isoform does not interfere with Myc-dependent cell cycle progression. Together, these data demonstrate that despite their essential role in nucleotide production, the cumulative activity of both PRPS enzymes is not required to carry out the global regulation of anabolic processes downstream of Myc hyperactivation. Rather, our data supports a model whereby increased PRPP production downstream of Myc overexpression in PRPS1 KO or PRPS2 KO cells is maintained by adjusting the rate of upstream oxPPP flux, thus explaining the increased concentration of NADPH in PRPS2 KO cells. Our data also indicates that PRPS activity is intrinsically linked to viability and apoptosis despite being uncoupled from Myc-dependent cell cycle progression and growth, suggesting that the perturbations that emerge upon PRPS KO are very specific metabolic effects linked to redox homeostasis.

**Figure 4.**
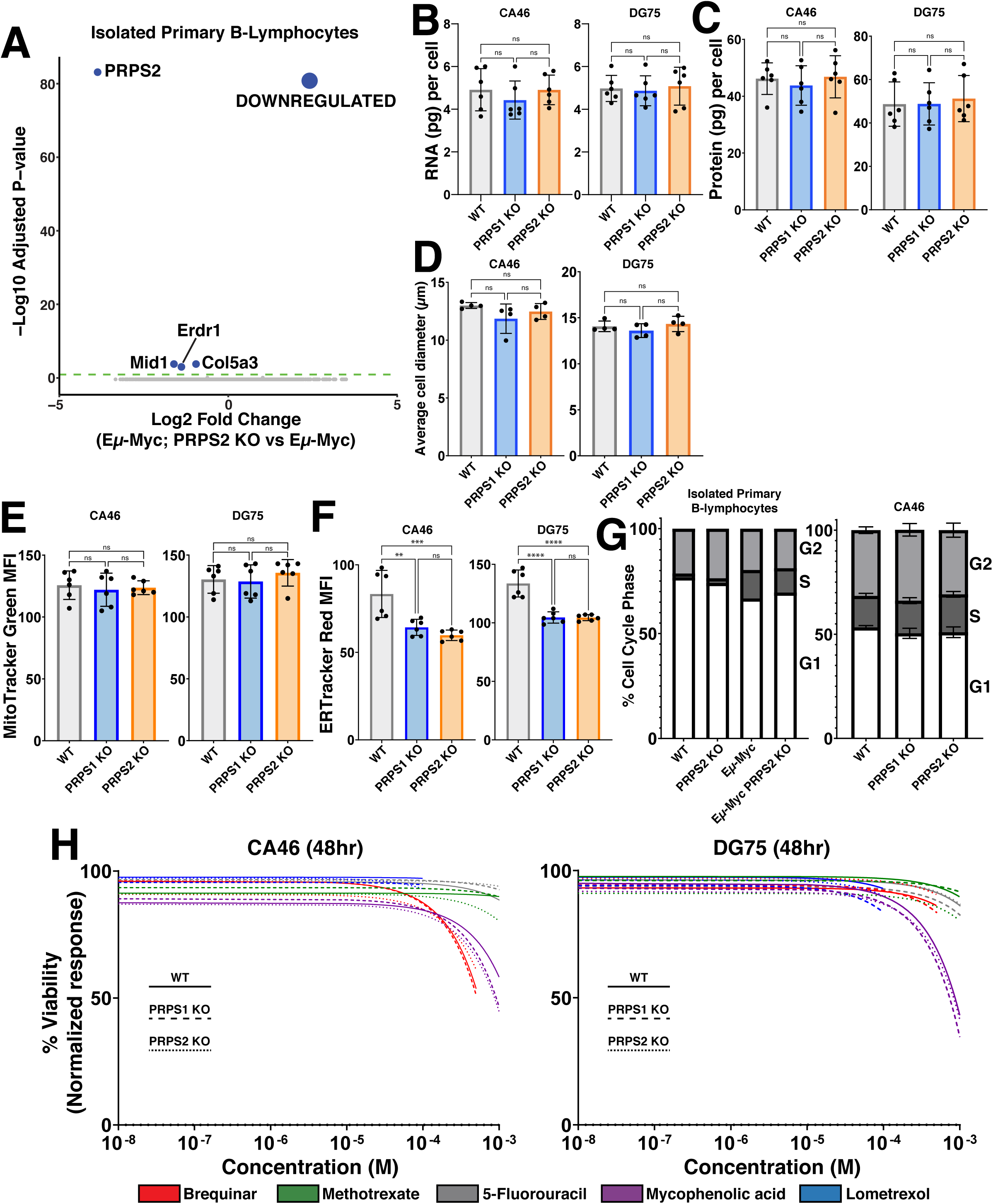
PRPS activity does not influence anabolic processes or cell cycle. (A) Volcano plot illustrating differentially expressed genes in Eµ-Myc; PRPS2 KO vs Eµ-Myc murine primary B lymphocyte cells (Statistical analysis detailed in Methods; dashed green line on volcano plot demarks significance of p<0.05). (B) RNA (pg) content per cell, (C) protein (pg) content per cell and (D) average cell diameter (µm) in WT, PRPS1- and PRPS2-KO cells of CA46 (left) and DG75 (right) cell lines. (E) Mitochondrial mass, measured via MitoTracker Green mean fluorescence intensity (MFI) and (F) ER expansion, measured via ER Tracker Red MFI of WT, PRPS1- and PRPS2- KO cells of CA46 (left) and DG75 (right) cell lines. (G) Cell cycle analysis profiling the percentage of cells in G1, S, and G2 phases for WT, PRPS2 KO, Eµ-Myc and Eµ-Myc; PRPS2 KO cells of murine primary B lymphocytes (left) and WT, PRPS1- and PRPS2- KO cells of CA46 cells (right). (H) Dose-response curves illustrating viability response to treatment with 5-fluorouracil, methotrexate, lometrexol, mycophenolic acid and brequinar in WT, PRPS1- and PRPS2- KO cells of CA46 (left) and DG75 (right) cell lines, normalized to vehicle treatment. X-axis represents the logarithmic scale of increasing drug concentration, Y-axis represents the normalized response as a viability percentage. Data was collected 48hrs post-treatment (Data represented as a mean of the normalized viability response of individual replicates at each concentration tested). For all panels, statistical analysis performed via One-Way ANOVA, bars represent mean ± s.d.; **p<0.01, ***p<0.001, ****p<0.0001, ns: not significant.

### Altering PRPP production does not induce sensitivity to inhibitors of nucleotide metabolism

To determine whether altered rates of PRPP production may have a threshold effect to induce sensitivity in Myc-overexpressing lymphomas to inhibitors of downstream PRPP-utilizing pathways, we treated our WT, PRPS1 KO and PRPS2 KO lymphoma cells with the following inhibitors: thymidine nucleotide production (5-fluorouracil; TYMS inhibitor) (Supplementary Fig. 4C), *de novo* pyrimidine biosynthesis (brequinar; DHODH inhibitor) (Supplementary Fig. 4D), *de novo* purine biosynthesis (lometrexol; GART inhibitor) (Supplementary Fig. 4E), folate metabolism (methotrexate; DHFR inhibitor) (Supplementary Fig. 4F), and guanosine nucleotide production (mycophenolic acid; IMPDH inhibitor) (Supplementary Fig. 4G). Not only did these compounds fail to elicit differential responses between WT and PRPS1 KO or PRPS2 KO cells, but they barely impacted viability at the highest concentrations tested. Indeed, only mycophenolic acid treatment achieved a 50% decrease in viability, which required millimolar concentrations of the drug (Fig. 4H). These findings that altering PRPP production via PRPS1 KO or PRPS2 KO does not induce sensitivity to inhibitors of nucleotide metabolism could perhaps be explained by a combination of decreased nucleotide catabolism and increased nucleotide salvage. These data demonstrate that targeting a single pathway of PRPP utilization is therapeutically insufficient, even in the context of PRPS1 KO or PRPS2 KO, reinforcing the concept that Myc-overexpressing cell viability is linked to redox homeostasis. These results also emphasize the importance of sustaining PRPP production, and demonstrate how Myc-overexpressing lymphomas adjust flux to accommodate more or less efficient PRPS complex configurations. These findings also underscore the difficulty in targeting the more aggressive, therapy-refractory tumors characterized by Myc hyperactivation, which exhibit tremendous metabolic flexibility by virtue of the myriad deregulated pathways that provide redundancy and resiliency^43^.

### PRPS-isozyme specific regulation reveals vulnerabilities in thioredoxin, glutathione redox pathways

To identify metabolic pathways and targets that elicit significant and differential responses between WT and PRPS KO lymphoma cells, we curated a library of over 100 different compounds with mechanisms of action involving metabolic processes (antioxidant properties, protein homeostasis, transcription, heme metabolism, lipid metabolism, glucose metabolism and apoptosis) and specific classes of enzymes (dehydrogenases, kinases and transporters) to pinpoint which Myc-dysregulated metabolic pathways could be leveraged as a combinatorial therapeutic approach (Supplementary Table 1). Each compound was tested at increasing concentrations in WT, PRPS1 KO and PRPS2 KO CA46 lymphoma cells such that normalized viability responses could be reported as an EC_50_ value. The results of this screening are displayed in Fig. 5A, with the response to treatment of each compound represented as ΔEC_50_ magnitude vs ΔEC_50_ significance in PRPS1 KO and PRPS2 KO cells. Of all the compounds tested in the CA46 lymphoma cell line, we found 7 that yielded both statistically significant and opposing sensitivities between PRPS1 KO and PRPS2 KO cells compared to WT cells, 5 of which recapitulated their behavior in DG75 cells: dithiothreitol (DTT), N-acetylcysteine (NAC), G6PDi-1, carmustine and auranofin. Of these 5 compounds, DTT and NAC both function as reducing agents^44,45^ and both sensitized PRPS2 KO cell viability compared to WT controls, while PRPS1 KO cells were more protected against cell death upon their treatment. This data suggests that DTT and NAC are more effective against Myc-overexpressing PRPS2 KO lymphoma cells because they exacerbate reductive stress^46^. Conversely, we found G6PDi-1, carmustine and auranofin all to be more toxic to PRPS1 KO cells compared to WT controls, whereas PRPS2 KO cells were more protected against cell death upon treatment. These compounds all create a more oxidative intracellular environment; G6PDi-1 suppresses NADPH production via blocking the first step in the oxPPP^27^, carmustine prevents the conversion of GSSG to GSH via carbamoylation of GSR^47^, and auranofin works in a similar fashion by preventing the reduction of thioredoxin disulfide bonds via inhibition of TRXR^48^. Importantly, the NADPH produced via G6PD is the reducing equivalent required to drive the reactions catalyzed by both GSR and TRXR, indicating that these pathways are intertwined to suppress oxidative stress (Fig. 5B).

**Figure 5.**
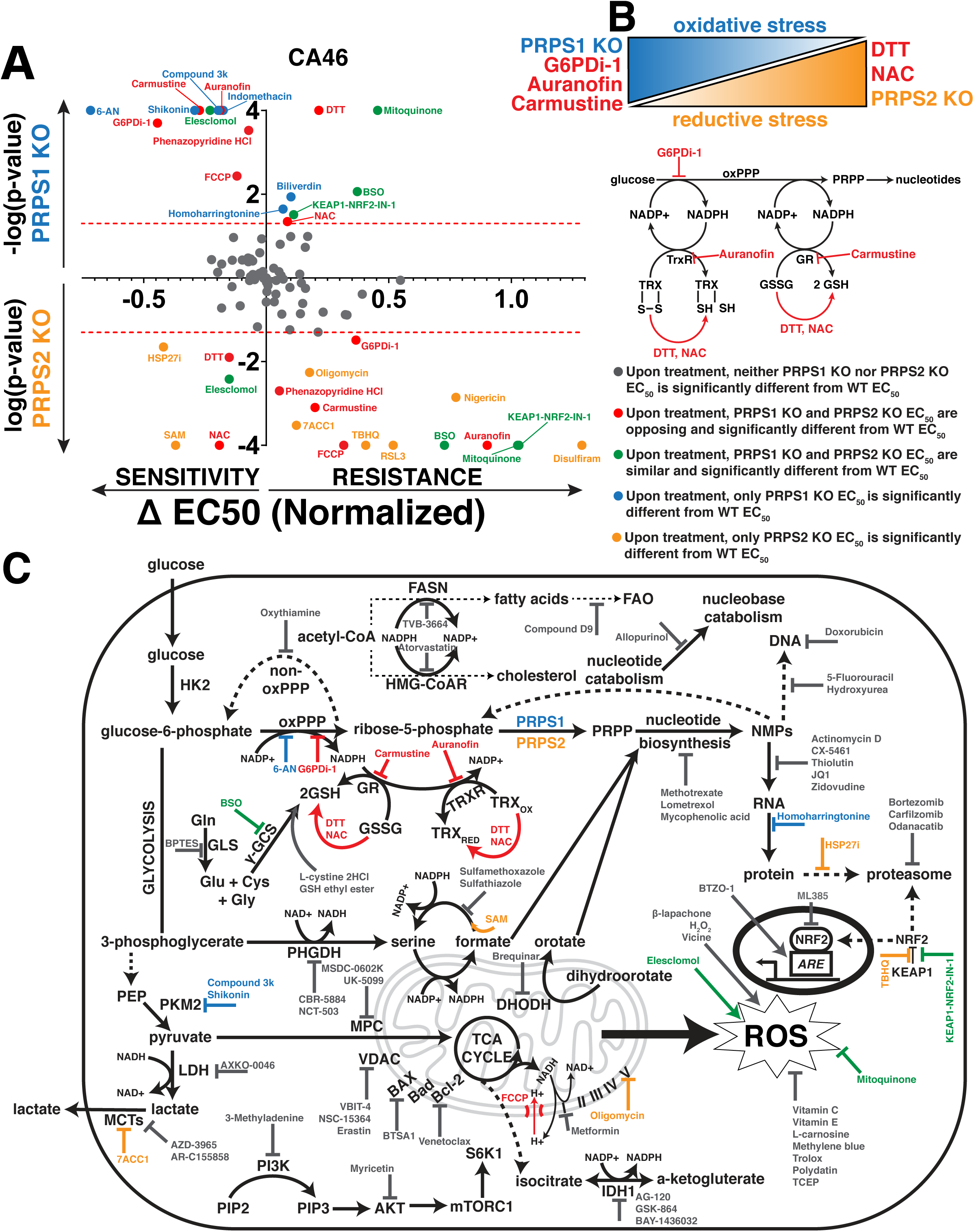
PRPS-isozyme specific augmentation of redox stress as a therapy. (A) Plot illustrating viability response of PRPS1- and PRPS2- KO CA46 cells to treatment with pharmacologic agents, normalized to vehicle treatment. X-axis represents magnitude of EC_50_ change, relative to WT EC_50_; right side of Y-axis represents resistance to compounds, left side of Y-axis represents sensitivity to compounds. Y-axis represents significance of EC_50_ change plotted as −log(p-value) or log(p-value), compared to WT EC_50_; PRPS1 KO cells represented above the X-axis, PRPS2 KO cells represented below the X-axis. Significance illustrated by dotted red lines on either side of the X-axis (p<0.05) (EC_50_ and significance determined via nonlinear regression model, fitting a variable slope for (log[x] vs normalized viability response)). (B) Schema illustrating the combinatorial approach used to generate oxidative stress and reductive stress in CA46 and DG75 lymphoma cells (top). Schema representing of the connection between oxPPP-derived NADPH and the thioredoxin and glutathione antioxidant pathways, with compounds highlighted in red (G6PDi-1, auranofin, carmustine, DTT, NAC) that elicited opposing and significantly differential sensitivities between PRPS1- and PRPS2- KO in CA46 and DG75 lymphoma cell lines. (C) Schematic representation of functional cooperativity map resulting from pharmacologic screening in CA46 cells. Color-coded legend provides distinctions for PRPS1- or PRPS2- KO EC_50_, compared to WT EC_50_.

The results from the screening are schematized in a functional cooperativity map (Fig. 5C), which represents a majority of the compounds tested and pathways affected based on the response or lack thereof in PRPS1 KO or PRPS2 KO CA46 cells compared to WT CA46 cells to establish key takeaways. Inhibiting NAD+/NADH-dependent dehydrogenase activity of enzymes such as IMPDH, LDH, phosphoglycerate dehydrogenase (PHGDH) and isocitrate dehydrogenase (IDH) resulted in no significant ΔEC_50_ between PRPS1 KO, PRPS2 KO or WT cells. The significant and differential responses upon inhibition G6PD, GSR and TRXR seem to be restricted to the glutathione and thioredoxin antioxidant systems, as inhibiting other NADP+/NADPH-dependent processes such as fatty acid synthase (FASN) and hydroxymethylglutaryl-coenzyme A reductase (HMG-CoAR) did not elicit differential responses upon treatment. These results are also specific to the NADPH-dependent reductive activity of the enzymes and not the presence of the antioxidant molecule GSH, as evidenced by a lack of differential response upon treatment with BSO, GSH ethyl ester or L-cystine 2HCl. Though many ROS-modulating compounds were tested, only NAC and DTT were found to generate significant differential sensitivities upon treatment in both cell lines. NRF2 activation or inhibition both failed to create differential sensitivities between KO cells, which aligns with our data suggesting that the KEAP1-NRF2 axis is not the primary effector for Myc-driven oxidative metabolism (Supplementary Fig. 1N). These data suggest that the effects revealed upon treatment with redox-modifying compounds are not a general phenomenon, but rather a very selective response. Also in line with our data (Fig. 4H), we observed that compounds interfering with different nodes of anabolic metabolism such as nucleotide metabolism, RNA synthesis and protein homeostasis did not reveal any significant and differential sensitivities between PRPS1 KO and PRPS2 KO cells, compared to WT cells. Additionally, no significant differential sensitivities were discovered when treating with inhibitors of pro- or anti-apoptotic proteins, mitochondrial membrane-bound transporters or mammalian target of rapamycin complex 1 (mTORC1) signaling. These data nominate the PRPS1 isozyme as a completely new potential therapeutic target in Myc-overexpressing lymphomas, as PRPS1 KO cells are sensitive to oxidative stressors that specifically interfere with the NADPH-dependent reductive activity of the thioredoxin and glutathione pathways. They also nominate PRPS2 ablation as a genetic tool to leverage alone or in combination with other reducing agents to generate toxic levels of reductive stress.

### Tuning PRPS activity as a therapeutic approach in Myc-overexpressing lymphoma

To test our model that PRPS1-dependent feedback sensing is a throttle on PRPS activity, thereby stifling oxPPP flux in lymphocytes, we assessed the ability of PRPS1 hypomorphic (A87T), superactive (D52H, H193L) and PRPS2 catalytically inactive (E39A) variants to rescue the differential sensitivities upon treatment with the 5 compounds that emerged from our screening. We observed that exogenous overexpression of PRPS2 and the superactive PRPS1 D52H and H193L mutants largely, if not completely, re-sensitized PRPS2 KO lymphoma cells to treatment with auranofin (Supplementary Fig. 5A), carmustine (Supplementary Fig. 5B) and G6PDi-1 (Supplementary Fig. 5C), while exogenous overexpression of PRPS1, hypomorphic PRPS1 A87T and catalytically inactive PRPS2 E39A failed to do so. In contrast, exogenous PRPS2 overexpression and superactive PRPS1 D52H and H193L mutants largely, if not completely, rescued the induced sensitivity of PRPS2 KO to treatment with DTT (Supplementary Fig. 5D) and NAC (Supplementary Fig. 5E), while exogenous overexpression of PRPS1, hypomorphic PRPS1 A87T and catalytically inactive PRPS2 E39A failed to do so. To rule out alternative mechanisms of toxicity linked to changes in redox, we assayed our cell lines for lipid peroxidation (Supplementary Fig. 5F), labile iron accumulation (Supplementary Fig. 5G) and global protein oxidation (Supplementary Fig. 5H), which failed to demonstrate a consistent and significant response between WT and PRPS KO lymphoma cells. Taken together, these data imply an exquisite specificity to the thioredoxin and glutathione networks within the Myc-regulated redox program comprised of the oxPPP-purine metabolism-OXPHOS circuit.

## Discussion

It’s been known for decades that Myc stimulates mitochondrial respiration and oxidative metabolism, with recent studies leveraging this knowledge to demonstrate the therapeutic benefit of attacking this core feature of Myc’s metabolic program^15–17^. Here, we unravel the mechanistic circuitry underlying the altered oxidative homeostasis, and in the process identify an important connection between cytosolic and mitochondrial redox metabolism. As lymphocytes are among the most rapidly proliferating cells in the body, replicating as quickly as 9 hours after the initial cell division^49^, *c-Myc* has also been shown to play an indispensable role in normal B cell activation and proliferation as a transcriptional amplifier with a core function in regulating the rate of nucleotide production^50^. Recent studies have also implicated oxidative metabolism as an essential component of B cell development by promoting selection, maturation and class-switch recombination of B cells^51,52^. Our data lends mechanistic support to these studies, suggesting that Myc engages OXPHOS machinery as one of the earliest metabolic alterations to exacerbate oxidative metabolism in lymphoma cells, simultaneous with that of nucleotide biosynthesis and transcription. Our work may also help explain observations regarding genetic disorders of the immune compartment such as severe combined immunodeficiency (SCID)^53^, Arts syndrome^54^, Charcot-Marie-Tooth disease-5 (CMTX5)^55^, DFN2^42^ and lupus erythematosus^56^ some of which are X-linked disorders arising from mutations in PRPS1 or PRPS2. Additionally, we provide mechanistic insight to help explain the immunosuppressive effects of pharmacological agents that target lymphocytes such as mycophenolic acid^57^ and mizoribine^58^ (IMPDH), allopurinol (XOR)^59^ and auranofin (TRXR)^60^. Our work functionally connects primary redox circuitries and helps explain these genetic and pharmacological observations, magnifying the critical balance of purine production and catabolism to maintain oxidative metabolism and overall immune competency of lymphocytes.

Our data demonstrates that supraphysiological levels of Myc activation generates oxidative stress, which has been shown to promote pro-tumorigenic processes such as proliferation, migration, angiogenesis, drug resistance and genomic instability in different cancers^13^. We show that induction of the oxPPP is concomitant with that of OXPHOS and serves a dual role in producing nucleotides and combatting ROS to sustain TRX and GSH. Our data is consistent with a model whereby the PPP functions as a cycle, its flux principally governed by the PRPS enzymes which serve as exit valves. Importantly, the PRPS enzymes and regulatory PRPSAPs exist in a dynamic megadalton multimeric assembly in lymphocytes that catalyzes the conversion of R5P to PRPP. Our group has established an evolutionary timeline for the origin of PRPS complex members and derived a model of PRPS complex assembly, where we discovered that the PRPS2 isoform arose from a gene duplication event where it diverged from and lost some of the allosteric feedback sensitivity of the ancestral PRPS1 isoform^34^. Here, we establish PRPS1 KO cells have lost the evolutionarily conserved allosteric brakes from the PRPS complex, permitting enhanced flux through the oxPPP to foster PRPS2-dependent nucleotide production. In PRPS2 KO cells however, these allosteric brakes are preserved, forcing enhanced cycling through the oxPPP which increases NADPH production as a byproduct of sustaining sufficient levels of PRPP and downstream nucleotides. Furthermore, we show that Myc-dependent upregulation of PRPS2 alters overall PRPS complex configuration, favoring a smaller dimeric assembly between PRPS1 and PRPS2 as opposed to the larger molecular weight complex that has been identified in Myc-low cells. These findings combine to suggest that the PRPS1:PRPS2 ratio is tunable to regulate overall PRPS activity based on the cell’s metabolic needs, with our data demonstrating that Myc overexpression favors an increase in PRPS2:PRPS1 to increase PRPS activity via the purine feedback insensitive PRPS2 isoform to enhance purine cycling and mitochondrial respiration.

The results of these findings nominate the PRPS isozymes as critical regulators of redox homeostasis in Myc-overexpressing lymphomas, which is uncoupled from Myc-dependent regulation of transcription, cell cycle control or other anabolic processes. We nominate a unique combinatorial therapeutic approach to take advantage of the Myc-dependent oxidative metabolism required for proliferating B cells, leveraging PRPS isoform-specific alterations in redox homeostasis to exacerbate vulnerabilities. Specifically, we discover that oxPPP-derived NADPH is necessary to drive the reactions catalyzed by TRXR and GSR, identifying inhibition of the PRPS1 isoform as a new potential therapeutic angle to sensitize cells to toxic levels of oxidative stress in these lymphomas. Conversely, while triggers of reductive stress-mediated cell death remain elusive due to a lack of appropriate markers and readouts^46^, we uncovered one of the few known loss-of-function genetic approaches to induce reductive stress (PRPS2 KO) and outlined a therapeutic approach combining reducing agents with PRPS2 KO to create toxic levels of reductive stress. Overall, this study highlights PRPS activity as the molecular rheostat dictating the redox state in Myc-driven lymphomas and provides a proof-of-concept in how the tunability of a metabolic enzyme may be exploited for therapeutic benefit in Myc-driven lymphoma.

## Materials & Methods

### Cell Culture

CA46 (ATCC #CRL-1648), DG75 (ATCC #CRL-2625), and P493-6 (Sigma-Aldrich #SCC279) cells were cultured in complete RPMI 1640 medium (Corning #10-041-CV) supplemented with 10% (v/v) fetal bovine serum (FBS) (Gibco #A5670701) and 1% (v/v) penicillin/streptomycin (Gibco #15140-122). P493-6 cells were cultured in the presence of 0.1 µg/mL tetracycline (Selleckchem #S4490) for 48hrs to achieve *c-Myc* suppression. For experiments involving the expression of Alternative Oxidase (AOX), cells were treated with 100 ng/mL doxycycline for 24hrs to induce expression. HEK293T cells (ATCC #CRL-3216) used for viral production were cultured in complete DMEM medium (Corning #10-013-CV) supplemented with 10% (v/v) FBS and 1% (v/v) penicillin/streptomycin. All cells were cultured at 37°C in a 5% CO_2_ incubator.

### Plasmid Generation

Lentiviral expression plasmids for shRNA-mediated knockdown were derived, as follows. shRNA sequences targeting *PRPS1* and *PRPS2* genes were created and ordered from IDT, as well as a non-targeting control sequence (Supplementary Table 2). Primers were annealed and subsequently ligated into the pLKO.1_puro cloning vector (Addgene #8453) via restriction cloning, using AgeI and EcoRI restriction sites. Lentiviral expression plasmids for ALFA-tagged peptides were derived, as follows. *Prps1* cDNA (Horizon Discovery #MMM1013-202859297) and *Prps2* cDNA (previously described^33^) were subcloned into the pMSCV_puro vector using the SalI and EcoRI restriction sites. The blasticidin expression cassette from pWZL_Blast_myc (Addgene #10674) was subcloned into FUGW_bleo (Addgene #14883) to replace bleomycin. The cDNAs encoding *PRPS1* and *PRPS2* were PCR amplified from the pMSCV-puro vector and ligated with the PCR-amplified FUGW_blasticidin backbone using Gibson assembly. Sequences encoding the ALFA epitope tag were engineered in the primers to C-terminally tag the proteins. Site-directed mutagenesis (SDM) was performed to introduce point mutations into ALFA-tagged PRPS1 and PRPS2 expression plasmids. To accomplish this, complementary primers containing the appropriate base substitutions were designed for the target gene and ordered from IDT (Supplementary Table 2). Template DNA was mixed with appropriate forward and reverse primers, along with Q5 Hot Start Hi-Fi 2X MM (NEB #M0494L), and the mixture was PCR amplified according to manufacturer’s instructions. KLD treatment was then performed on the PCR product using 10X KLD enzyme mix (NEB #M0554S) and 2X KLD reaction buffer (NEB #B0554A) to circularize the mutated plasmid. DNA product was then transformed using 5-alpha Competent *E. coli* cells (NEB #C3040I), and subsequently isolated via GeneJET Plasmid MiniPrep Kit (Thermo Scientific #K0503), according to manufacturer’s protocol. Sanger sequencing was performed to confirm incorporation of mutation. Lentiviral expression plasmids for TPNOX and mitoTPNOX were derived, as follows. TPNOX and mitoTPNOX cDNA were amplified from Addgene plasmids #87853 and #87854, respectively, with primers containing sequences encoding the ALFA epitope tag to C-terminally tag the proteins. Subsequently, the constructs were cloned into the aforementioned FUGW_blasticidin plasmid assembly via restriction digest using AgeI and EcoRI restriction sites. Lentiviral expression plasmid for doxycycline-inducible mammalian AOX expression was obtained from Addgene (Addgene #177984). Retroviral expression plasmid for mammalian NDI1 expression was obtained from Addgene (Addgene #72876).

### Virus Production and Transduction

Lentivirus was produced in HEK293T cells by co-transfecting the lentiviral transfer vector with psPAX2 (Addgene #12260) and pMD2.G (Addgene #12259). Retrovirus was produced in HEK293T cells by co-transfecting the retroviral transfer vector with pUMVC (Addgene #8449) and pMD2.G. Transfection was carried out using PolyFect transfection reagent (QIAGEN #301105), and the media was replaced after 24 hours. Viral supernatant was collected at 48 hours and 72 hours post-transfection, filtered (45µm), and concentrated using Lenti-X (Takara #631232) or Retro-X (Takara #631456) concentrators.

Viral transduction of suspension cell culture was achieved via spinfection, as follows. 3×10^6^ cells were collected, per transduction, and resuspended in 1mL of filtered and concentrated viral media containing 8µg/mL polybrene (Millipore #TR-1003-G) to enhance transduction efficiency. Viral cell suspensions were centrifuged in 24-well plates at 200xg for 1hr, before subsequent incubation at 37°C. Complete media was added to the viral media (1:1) overnight, and the process was repeated 1-2X in consecutive days. After final spinfection was completed, cells were left to recover in complete media for 24hrs prior to the addition of the appropriate selection media.

### CRISPR-Cas9 Genome Editing

*PRPS1* and *PRPS2* knockouts in CA46 and DG75 cell lines were achieved using a CRISPR-Cas9 nickase system, as follows. BfuAI restriction sites, tracrRNA, and a GFP tag were inserted into a pLKO.1_TRC cloning vector (Addgene #10878). Complementary PRPS1 and PRPS2 crRNA oligonucleotides were generated for both the top (+) and bottom (-) strands (Supplementary Table 2) of the respective target gene to facilitate Cas9 D10A-mediated nickase activity. Annealed crRNA oligonucleotide strands were inserted into the BfuAI-digested pLKO.1 backbone. The (+) sgRNA and (-) sgRNA pLKO plasmids were then electroporated with Cas9_D10A Nickase (Addgene #41816) at a 1:1:1 ratio via the Invitrogen Neon Transfection System, according to manufacturer’s protocols. After 48 hours, cells were single-cell sorted into individual wells of a 96-well plate based on GFP expression, via the BD FACSAria. Single cells were subject to clonal expansion until sufficient material was able to be gathered for thorough knockout validation via Western Blotting, PCR, and DNA sequencing.

### SDS-PAGE and Western Blotting

For denaturing cell lysis, cells were first washed 1X with ice-cold PBS (Corning #21-040-CV) and subsequently lysed using RIPA buffer (Thermo Scientific #89901) supplemented with 1X protease and phosphatase inhibitor cocktail (Thermo Scientific #78446). Cleared protein lysates were quantified using BCA reagent (Thermo Scientific #23227), and subsequently combined with 1X Laemmeli sample buffer prior to separation on 10% TGX Fastcast gels (BioRad #1610173) in 1X Tris/Glycine/SDS running buffer (Thermo Scientific #PI28362) at 120V for ∼1hr. Separated proteins were then transferred onto 0.2µm PVDF membranes (BioRad #1704156) using the Bio-Rad Trans-Blot Turbo Transfer system. PVDF membranes were subsequently blocked for 1hr at room temperature (RT) with 5% (w/v) milk in Tris-Buffered Saline containing 0.1% Tween-20 (Thermo Scientific #J20605.AP) (TBS-T). After blocking, the membranes were washed (3X in TBS-T) and incubated overnight at 4°C with primary antibodies (diluted 1:1000) prepared in 3% BSA (Thermo Scientific #J64100.22) in TBS-T. Membranes were again washed (3X in TBS-T) and incubated at RT with corresponding secondary antibodies from Jackson ImmunoResearch (diluted 1:25000) in 5% (w/v) milk in TBS-T. Blots were visualized using chemiluminescent substrates from Thermo Scientific via the Bio-Rad ChemiDoc Touch Imaging System. To facilitate re-probing, Restore PLUS Western Blot Stripping Buffer (Thermo Scientific #46430) was used to strip the blots after imaging. To determine oxidative state of proteins, cells were first washed 1X with ice cold PBS and subsequently incubated in 1% CHAPS detergent (Millipore #75621-03-3) containing 1 mmol/L PEG-PCMal (Dojindo #SB20) for 30 min at 37°C. Cleared protein lysates were quantified using BCA reagent and subsequently resolved via SDS-PAGE in non-reducing conditions (DTT-free 1X Laemmeli sample buffer, no boil).

### Antibodies

Primary antibodies used were: c-Myc (Cell Signaling #9402), ACLY (Cell Signaling #13390), ATIC (Abcam #ab33520), RRM2 (Novus Biologicals #NBP1-31661), eIF4E (Cell Signaling #2067), NPM (Cell Signaling #3542), NDUFA9 (Abcam #ab14713), SDHA (Cell Signaling #11998), UQCRC1 (Invitrogen #459140), COXIV (Abcam #ab202554), ATP5A (Abcam #ab14748), BiP (Cell Signaling #3177), Cleaved PARP-1 (Abcam #32064), β-Actin (Cell Signaling #4970; Cell Signaling #3700), RPB1 CTD (phospho-S2) (Cell Signaling #13499), RPB1 CTD (phospho-S5) (Cell Signaling #13523), RPB1 CTD (phospho-T4) (Cell Signaling #26319), RPB1 CTD (Cell Signaling #2629), POLR1A (Cell Signaling #24799), POLR3A (Cell Signaling #12825), H3K27Ac (Cell Signaling #4353), RPE (Abcam #ab128891), 6PGD (Abcam #ab129199), PGLS (Abcam #ab135771), HK2 (Cell Signaling #2867), RPIA (Santa Cruz #sc-515328), PRPS1/2/3 (Santa Cruz #sc-376440), G6PD (Santa Cruz #sc-373886), KEAP1 (Cell Signaling #8047), HO-1 (Cell Signaling #43966), NQO1 (Santa Cruz #sc-32793), TKT (Santa Cruz #sc-390179), PRPS1/2 (Santa Cruz #sc-100822), Anti-Puromycin (Millipore #MABE343), GAPDH (Cell Signaling #5174), PRPS1 (Proteintech #15549-I-AP), PRPS2 (Sigma #SAB2107995), CAD (Cell Signaling #93925), PRPSAP1 (Santa Cruz #sc-398422), PRPSAP2 (Proteintech #17814-1-AP), TCP1-η (Santa Cruz #sc-271951), AK2 (Santa Cruz #sc-374095), ALFA-HRP (SynapticSystems #N1505-HRP), CTPS (Abcam #ab133743), UMPS (Santa Cruz #sc-398086), HPRT (Abcam #ab109021), α-Tubulin (Abcam #ab176560), AHCY (Sigma #HPA041225), RPS7 (Cell Signaling #sc-377317), RPL11 (Cell Signaling #18163), eIF2α (phospho-S51) (Cell Signaling #3398), eIF2α (Cell Signaling #5324), H6PD (GeneTex #GTX101500), SOD2 (Proteintech #24127-1-AP), TRX1 (Cell Signaling #2429), Ki67 (Abcam #ab16667), Cyclin D2 (Cell Signaling #3741), Cyclin E1 (Proteintech #11554-1-AP), Rb (phospho-S807/811) (Cell Signaling #8516), Rb (Cell Signaling #9309), Histone H3 (phospho-S10) (Cell Signaling #9701), Histone H3 (Abcam #ab176842), p21 (Santa Cruz #sc-6246), p27 (Cell Signaling #3688), Bim (Cell Signaling #2933), p62 (Cell Signaling #39749), TRX2 (Santa Cruz #sc-133201), Catalase (Cell Signaling #14097), XO (Abcam #ab109235), NNT (Santa Cruz #sc-390236), PRDX1 (Cell Signaling #8499), LDHA (Cell Signaling #3582), GCLM (Abcam #ab126704), GMPR2 (Invitrogen #PA5-67012), NDUFA5 (Proteintech #16640-1-AP), NDUFA12 (Proteintech #15793-1-AP), NDUFAF2 (Proteintech #13891-1-AP), NDUFAF4 (Proteintech #26003-1-AP), NDUFB5 (Cusabio #CSB-PA015652ESR1HU), NDUFS1 (Cell Signaling #60153), NDUFS2 (GeneTex #GTX114924), NDUFS3 (Proteintech #15066-1-AP), NDUFS4 (GeneTex #GTX105662), NDUFS6 (Proteintech #14417-1-AP), NDUFV2 (Proteintech #15301-1-AP), MTCO1 (Abcam #ab14705), AMPKα (phospho-T172) (Cell Signaling #2535), AMPKα (Cell Signaling #2532), IDH1 (Cell Signaling #3997), IDH2 (Cell Signaling #60322), MDH2 (Santa Cruz #sc-293474), HMGCR (Santa Cruz #sc-271595), NRF1 (Cell Signaling #46743), TRXR1 (Cell Signaling #15140), TRXR2 (Santa Cruz #sc-365714), GLRX (Proteintech #15804-1-AP), GLRX2 (Proteintech #13381-1-AP), GSR (Santa Cruz #sc-133245), Bak (Cell Signaling #12105), Bax (Cell Signaling #5023), Cleaved Caspase-3 (Cell Signaling #9661), FASN (Cell Signaling #C20G5), FLC (Santa Cruz #sc-390558), NDUFB7 (Proteintech # 14912-1-AP), NDUFB9 (Abcam #ab 200198), ECSIT (Sigma #HPA042979), IMPDH1 (Abcam #ab137112), IMPDH2 (Abcam #ab131158), MDH1 (Proteintech #15904-1-AP), MTHFD1L (Cell Signaling #14999), SAMHD1 (Origene #TA502157S), SHMT1 (Cell Signaling #80715), SHMT2 (Cell Signaling #12762), PHGDH (Santa Cruz #sc-100317), DHFR (Abcam #ab124814), FLAG (Cell Signaling #8146).

Secondary Antibodies used were: Anti-mouse (Jackson ImmunoResearch #115-035-003), Anti-rabbit (Jackson ImmunoResearch #111-035-003).

### Mice

C57BL/6 mice (JAX #000664) and Eμ-MYC mice^19^ (JAX #002728) were obtained from The Jackson Laboratory. PRPS2 KO mouse strain used for this research project was created from ES cell clone EPD0110_2_A05, obtained from KOMP Repository (www.komp.org) and generated by the Wellcome Trust Sanger Institute. Mice were generated at the Transgenic Animal and Genome Editing (TAGE) core at Cincinnati Children’s Hospital Medical Center (CCHMC).

### Primary B Lymphocyte Isolation

Post-euthanasia (CO_2_ asphyxiation followed by cervical dislocation), spleens of each mouse (6w) were dissected and immediately stored in ice-cold 1X PBS. Scalpel was used to break through the integrity of the spleen casing, prior to filtration through 40µm cell strainers (Corning #431750) using a 1mL syringe plunger and 1X PBS. Cell suspension was then centrifuged at 300xg for 5min, before the supernatant was discarded and pelleted cells were resuspended in 5mL 1X red blood cell (RBC) lysis buffer (BioLegend #420301), according to manufacturer’s protocol. Cell suspension was again centrifuged at 300xg for 5 min, repeating the RBC lysis process until cell pellets were homogenously white in color. 10mL PBS was added to neutralize the lysis buffer, and cell suspension was again centrifuged at 300xg for 5min. Cell pellets were resuspended in 1X PBS, counted using Trypan blue (Sigma #T8154) at 1/10 dilution via Countess II FL, and then prepared for B-cell isolation via negative selection (Miltenyi #130-090-862), according to manufacturer’s protocol. To induce B-cell activation, lipopolysaccharides (LPS, Invitrogen #00-4976-03) were added to complete RPMI 1640 media at a 1X concentration (5µg/mL) immediately following isolation.

### RNA Sequencing

Following isolation, RNA was isolated from primary B lymphocytes using Trizol reagent (Invitrogen #15596018), according to manufacturer’s instructions. RNA samples were processed by the University of Cincinnati Department of Environmental and Public Health Sciences Genomics, Epigenomics and Sequencing Core. NEBNext Ultra II Directional RNA Library Prep kit was used for RNA-seq library preparation. Sample RNA passed through a quality control step using Agilent RNA bioanalyzer QC and sequenced via PolyA RNA-seq (SR 1x85 bp, ∼25M reads, >20M pass filter) on Illumina NextSeq 550 sequencer^61^. The differentially expressed genes (DEGs) analysis was performed using DEseq2 with Illumina’s platform and the volcano plot was generated using ggplot2 in R (v4.3.3). DEGs were selected by limiting the adjusted p-value < 0.05. For comparison 4, from the total of 15,430 genes analyzed in our study, 3,785 were significantly upregulated and 3,749 were significantly downregulated. For comparison 5, from the total of 17,611 genes analyzed, only 4 of them were significantly downregulated and none of them are significantly upregulated.

### Oxygen Consumption Rate (OCR), Extracellular Acidification Rate (ECAR)

The Agilent Seahorse XF Cell Mito Stress Test (Agilent #103015-100) was used to measure OCR and ECAR in P493-6 cells that had been removed from tetracycline-containing media at the indicated time points. The day prior to the assay, Seahorse cell culture plates (Agilent #103799-100) were coated in poly-d-lysine (Gibco #A3890401) according to manufacturer’s protocol to ensure adherence of suspension cells. 10^5^ cells were plated in 50µL Seahorse XF RPMI media (Agilent #103576-100) [(supplemented with 1mM pyruvate (Agilent #103578-100), 2mM glutamine (Agilent #103579-100) and 10 mM glucose (Agilent #103577-100)] per replicate, based on previously determined optimal seeding density, and plates were centrifuged at 200xg for 1min to adhere cells. Cells were incubated at 37°C in a non-CO_2_ containing incubator for 30 min, prior to the addition of 130µL complete Seahorse XF RPMI media. Cells were incubated once more for an additional 30 min prior to running the assay. 1hr prior to running the assay, H_2_O added overnight to Seahorse FluxPak sensor cartridge (Aglient #103792-100) was replaced with XF Calibrant Solution (Agilent #100840-000) and incubated at 37°C in a non-CO_2_ containing incubator. Immediately prior to running the assay, sensor cartridge was loaded with compounds in the following orientation: 20µL oligomycin (15µM) to Port “A”; 22µL FCCP (10µM) to Port “B”; 25µL rotenone + antimycin A (5µM each), such that the final working concentrations of each compound were 1.5µM, 1µM and 500nM, respectively. Injections were sequentially added in accordance with the Seahorse XF Mito Stress Test using the Agilent WAVE software and Agilent XFe96 Seahorse Analyzer.

### AlamarBlue Intracellular Reduction Assay

Cells were seeded in black-walled, clear-bottom 96-well plates at a density of 2.5×10^5^ cells/well (DG75, CA46, P493-6) or 5×10^5^ cells/well (isolated primary B lymphocytes) in 180µL complete RPMI 1640 medium. 20µL AlamarBlue reagent (Thermo Scientific #DAL1100) was added to each well to achieve a final concentration of 10% (v/v), prior to incubation for 4hrs at 37°C in a 5% CO_2_ incubator. Post-incubation, AlamarBlue fluorescence was measured using the BMG Labtech CLARIOStar microplate reader, with ex/em of 560nm/590nm. Data was collected via the MARS data analysis software, with background fluorescence being subtracted from all experimental wells. For drug treatment assays involving P493-6 cells, cells were treated for 48hrs prior to assay at the following concentrations: vehicle control (DMSO)(1% v/v), chloramphenicol (100µM), G6PDi-1 (10µM), allopurinol (100µM) and brequinar (100µM).

### Fluorescent Dyes

For each fluorescent dye used, 1.5×10^6^ cells were collected per replicate, in triplicate. Cell suspensions were centrifuged and washed with 1X PBS prior to incubation with the respective dye, with the working concentration and incubation time for each dye as follows: MitoSOX Red Superoxide Indicator (Thermo Scientific #M36008)(100nM, 30 min), CM-H_2_DCFDA (Thermo Scientific #C6827)(500nM, 30 min), MitoTracker Green (Thermo Scientific #M7514)(100nM, 30 min), ER Tracker Red (Thermo Scientific #E34250)(1µM, 30 min), 2-NBDG (Thermo Scientific # N13195)(100µM, 15 min), LysoTracker Green (Thermo Scientific #L7526)(50nM, 30 min), FerroOrange (Dojindo #F374-10)(1µmol/L, 30 min), BODIPY 581/591 C11 (Thermo Scientific #D3861)(5µM, 30 min). After incubation, cell suspensions were collected, centrifuged and washed with 1X PBS. Cells were again centrifuged and resuspended in FACS buffer (2.5% FBS in PBS) on ice for subsequent FACS analysis via BD LSRFortessa

### Luciferase-based Metabolite GLO Assays

The following kits were utilized for luciferase-based GLO-assays and executed in accordance with manufacturer’s protocols: NAD+/NADH-Glo Assay (Promega #G9072); NADP+/NADPH-Glo Assay (Promega #G9082); GSH/GSSG-Glo Assay (Promega #V6612). All absorbance measurements were obtained via BMG Labtech CLARIOStar microplate reader, and data was collected via MARS data analysis software.

### Per Cell Measurements

5×10^6^ cells were collected per analysis (RNA, protein, cell size), in triplicate. Cells were collected, centrifuged and washed with 1X ice-cold PBS. Protein lysates were obtained from each cell line in accordance with SDS-PAGE/Western Blotting protocol, followed with subsequent quantification via BCA assay. RNA was isolated from cells via PureLink RNA Mini Kit (Thermo Scientific #12183018A), according to manufacturer’s protocol. RNA concentration was obtained via Qubit4 Fluorometer, using the Qubit RNA BR Assay Kit (Thermo Scientific #Q10210), according to manufacturer’s protocol. Average cell size was obtained with the Countess II FL, using trypan blue staining at a 1:1 ratio.

### Cell Cycle Analysis

For cell cycle analysis experiments, 5×10^5^ cells were analyzed per replicate, in triplicate. Cells were centrifuged and washed with 1X ice-cold PBS, prior to fixation in 1.2mL ice-cold 66% ethanol in PBS with gentle vortexing. Cells were stored at 4 degrees for at least 2 hours (stable up to 4w) prior to centrifugation and subsequent washing with 1X PBS. Cells were again centrifuged, supernatant was discarded, and cells were resuspended in 200µL PBS containing

Propidium Iodide (Thermo Scientific #P3566) at a final concentration of 50µg/mL and RNase (Roche #11119915001) at a final concentration of 50µg/mL. Cells were incubated in the dark at 37°C in a 5% CO_2_ incubator for 30 minutes prior to FACS analysis via BD LSRFortessa.

### Puromycylation Assay

For each puromycylation assay, 1×10^6^ cells were collected and plated in 6-well plates, per replicate. Puromycin (Gibco #A11138-03) was added at a working concentration of 10µg/mL to each replicate for 30 min, before cells were collected and prepared in accordance with the SDS-PAGE/Western Blotting protocol.

### Size Exclusion Chromatography

Cells were lysed using non-denaturing lysis buffer (50 mM Tris-Cl, pH 7.5, 200 mM NaCl, 1% digitonin, 1mM TCEP (tris(2-carboxyethyl)phosphine), 1mM MgCl2, benzonase and 1X protease and phosphatase inhibitor cocktail) for 20 mins on ice. The cell lysates were then clarified by centrifugation at 15,000 × g for 15 mins at 4 °C and subsequently filtered using a 0.22 µm filter. About 200 μg of cell lysates were loaded onto a Superose 6 Increase 3.2/300 column (GE Healthcare #29-0915-98) at the flow rate of 0.04 mL/min using Thermo Vanquish UHPLC, using mobile phase (0.1M Na_2_HPO_4_, 0.1M NaCl, pH 7.5). After passing through the void volume, the sample fractions were collected and concentrated using a 3K MWCO filter (Thermo #88512) and analyzed via SDS-PAGE/Western Blotting. Gel filtration calibration kits (GE #28-4038-41, GE #28-4038-42, Sigma #MWGF200-1KT) were used to monitor column performance over time. Different internal standards were probed for molecular weight calibration: CAD^62^ [(GLN-dependent carbamoyl phosphate synthetase (CPS-2), aspartate transcarbamylase (ATC), dihydroorotase(DHO)], TCP-1η^63^ (T-Complex Protein 1 subunit eta), FASN (Fatty Acid Synthase)^64^ and FLC (Ferritin Light Chain)^65^ form complexes of around 1500 kDa, 900 kDa, 540 kDa and 480 kDa, respectively, while HK2 (Hexokinase 2) and AK2 (Adenylate Kinase 2) are mostly monomeric at 102 kDa and 26 kDa, respectively. The elution profile of PRPS complex components was compared with that of CAD, TCP-1η, FASN, FLC, HK2 and AK2, which were used as internal standards to estimate the size of the PRPS enzyme complex. Internal standards were probed in every SEC run.

### Global Protein Oxidation

Global protein oxidation was determined via the OxyBlot Protein Oxidation Detection Kit (Sigma-Aldrich #S7150), according to manufacturer’s protocol, and detected via SDS-PAGE/Western blotting.

### Pharmacological Dose-Response Screening

For each dose-response curve, cells were plated at a density of 6.5×10^4^ in 180µL of complete RPMI 1640 media in individual wells of a black-walled, clear-bottom 96-well plate with 4 blank wells containing media with no cells. 20µL of AlamarBlue cell viability reagent was added to each well, such that the final AlamarBlue concentration was 10% (v/v). Cells were left to incubate for 4hrs at 37°C in a 5% CO_2_ incubator, prior to obtaining the initial 0hr reading using the BMG Labtech CLARIOStar microplate reader using ex/em of 560nm/590nm. Stock concentrations of each compound were prepared in increasing 100X increments, such that adding 2µL of each stock would yield the final desired working concentration. After the initial 0hr reading, 2µL of each prepared stock was added to each cell line, and subsequent microplate readings were obtained via the BMG Labtech CLARIOStar microplate reader at 24, 48, and 72hrs post-treatment. If sufficient concentrations were not used to yield a proper EC_50_ dose-response curve, the experiment was repeated with a more accurate range of concentrations (unless solubility of original stock concentration proved to be a limiting factor). All data was collected via the MARS data analysis software.

### Quantification and Statistical Analysis

Countess II FL was used in conjunction with trypan blue for all cell counting and analysis of cell size. MARS data analysis software was used for the collection of all data obtained via the BMG Labtech CLARIOStar microplate reader. FlowJo was used for the analysis of all (.fcs) files obtained via Flow Cytometry. GraphPad Prism was used for the normalization, statistical analysis, and plotting of all data sets. Agilent WAVE software was used to conduct and collect data from Seahorse experiments. Bio-Rad ChemiDoc Touch Imaging System and Bio-Rad ImageLab were used for the acquisition and representation of all chemiluminescent Western Blot images. Adobe Illustrator was used to compile and generate all figures in their final form. Unless otherwise specified, n≥3 replicates were used per experiment, with One-Way ANOVA statistical test used to determine significance for all quantitative data except for dose-response curves, where a nonlinear regression model fitting a variable slope for (log[x] vs normalized viability response) was used to determine EC_50_ and significance.

## Acknowledgements

We thank the Transgenic Animal and Genome Editing (TAGE) core at the Cincinnati Children’s Hospital Medical Center (CCHMC) for the generation of the PRPS2 KO mouse strain used for this research project, which was created from ES cell clone EPD0110_2_A05 obtained from KOMP Repository (www.komp.org) and generated by the Wellcome Trust Sanger Institute. We thank the Genomics, Epigenetics and Sequencing (GES) core at the University of Cincinnati for conducting RNA sequencing. We thank B. Ehmer for assistance with the training, maintenance and oversight of Flow Cytometry equipment in the Advanced Cell Analysis Service Center (ACASC) at the University of Cincinnati. We thank K. Patra, C. Andreani and C. Bartolacci for their insights during the manuscript review process.

## Author contributions

Conceptualization: ACM, JTC

Methodology: ACM, JTC

Investigation: ACM, JTC, BK, SZ, KRG, JGP

Formal analysis: ACM, JY

Visualization: ACM, JTC, JY

Funding acquisition: JTC, MCK

Project administration: JTC

Supervision: JTC, MCK, JM

Writing – original draft: ACM, JTC

Writing – review & editing: ACM, JTC, BK, KRG

## Data and materials availability

All data supporting the findings of this study are available within the Article and its Supplementary Information. Raw and processed RNA sequencing data may be obtained via accession no. GSE282435 at https://www.ncbi.nlm.nih.gov/geo/query/acc.cgi?acc=GSE282435.

---

**Supplementary Figure 1 – Related to Figure 1.**
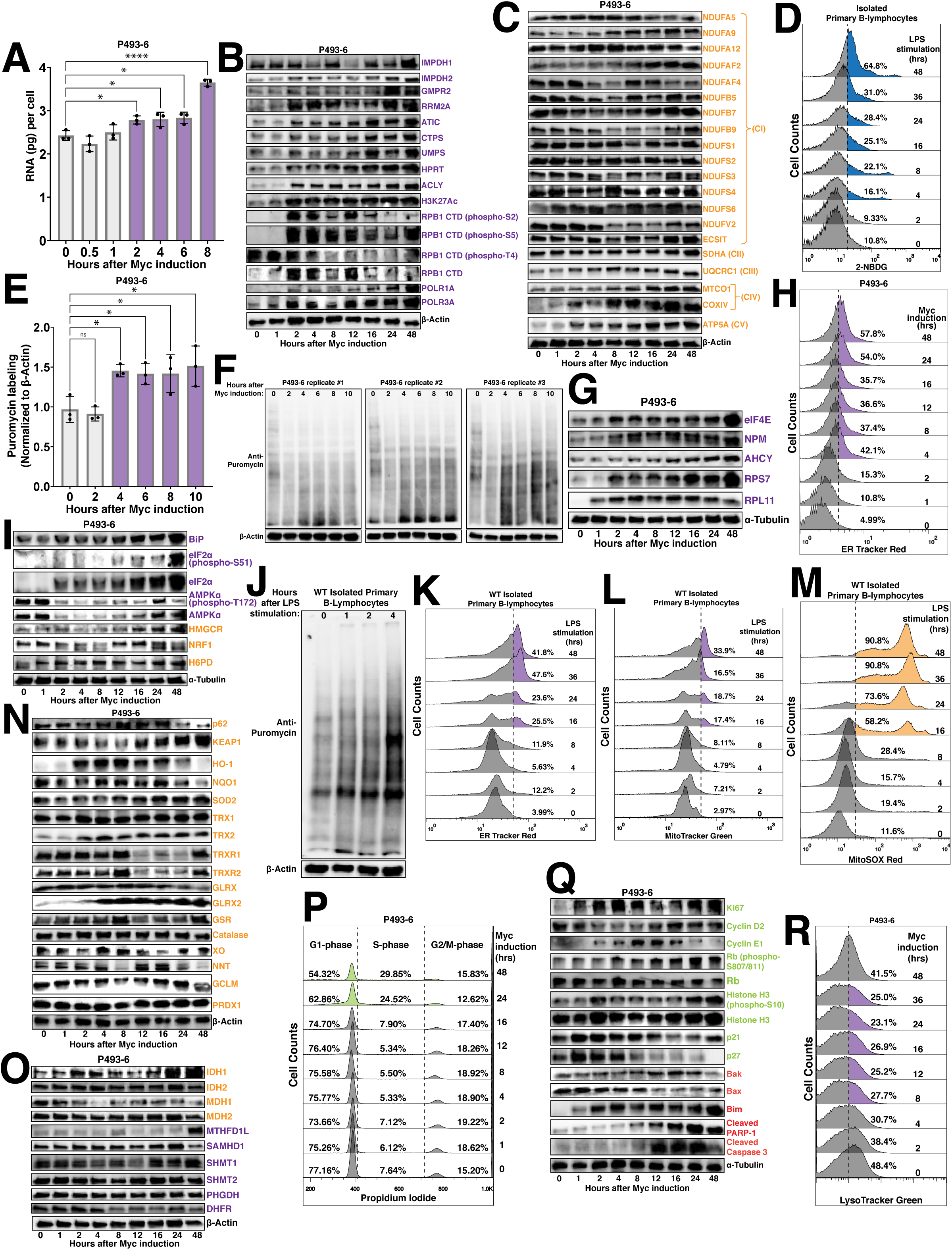
(A) RNA content per cell measured in P493-6 cells over an 8hr time course following tetracycline removal to induce Myc expression. Western blot of (B) nucleotide biosynthesis gene expression and transcriptional regulation and (B) nuclear-encoded mitochondrial complex gene expression in P493-6 cells over a 48hr time course following tetracycline removal to induce Myc expression. β-Actin used as a loading control. (C) Glucose uptake in WT murine primary B lymphocytes over a 48hr time course following LPS stimulation, measured via 2-NBDG. (E,F) Puromycylation assay measuring protein synthesis in P493-6 peptides over a 10hr time course following tetracycline removal to induce Myc expression, normalized to β-Actin. (G) Western blot of translational regulation in P493-6 cells over a 48hr time course following tetracycline removal to induce Myc expression. α-Tubulin used as a loading control. (H) Endoplasmic reticulum (ER) expansion in P493-6 cells over a 48hr time course following tetracycline removal to induce Myc expression, measured via ER Tracker Red. (I) Western blot of ER-localized and ER-stress response protein expression in P493-6 cells over a 48hr time course following tetracycline removal to induce Myc expression. α-Tubulin used as a loading control. (J) Puromycylation assay measuring protein synthesis in WT murine primary B lymphocyte peptides over a 4hr time course following LPS stimulation. β-Actin used as a loading control. (K) ER expansion, measured via ER Tracker Red and (L) mitochondrial mass, measured via MitoTracker Green in WT murine primary B lymphocytes over a 48hr time course following LPS stimulation. (M) Mitochondrial ROS accumulation in WT murine primary B lymphocytes over a 48hr time course following LPS stimulation, measured via MitoSOX Red. (N) Western blot analysis of antioxidant response element (ARE) metabolic enzyme expression in P493-6 cells over a 48hr time course following tetracycline removal to induce Myc expression. β-Actin used as a loading control. (O) Western blot analysis of tricarboxylic acid (TCA) cycle and folate metabolism enzyme expression in P493-6 cells over a 48hr time course following tetracycline removal to induce Myc expression. β-Actin used as a loading control. (P) Cell cycle analysis of P493-6 cells over a 48hr time course following tetracycline removal to induce Myc expression, measured via Propidium Iodide. Cell cycle profiling quantified as a percentage of cells in G1, S and G2/M phases at each time point. (Q) Western blot of cell cycle regulation and apoptosis in P493-6 cells over a 48hr time course following tetracycline removal to induce Myc expression. α-Tubulin used as a loading control. (R) Lysosomal content in P493-6 cells over a 48hr time course following tetracycline removal to induce Myc expression, measured via LysoTracker Green. For all panels, statistical analysis performed via One-Way ANOVA, bars represent mean ± s.d.; *p<0.05, ****p<0.0001, ns: not significant. For all histograms, upregulation/downregulation quantified as a percentage of the population to the right of the dashed line at each time point.

**Supplementary Figure 2 – Related to Figure 2 and Figure 3.**
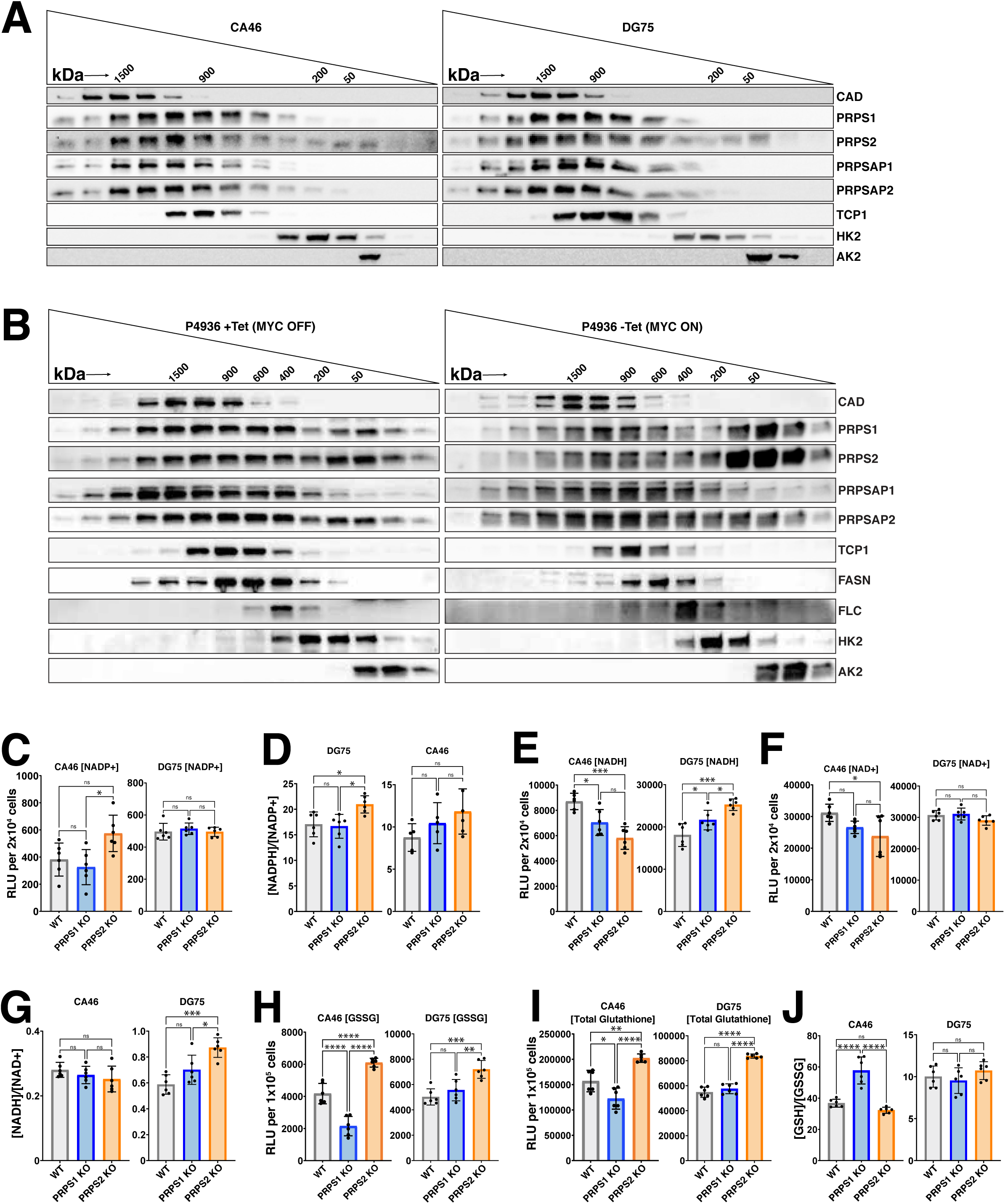
Western blot analysis of PRPS complex coordination using fractions collected from size exclusion chromatography (SEC) runs to visualize PRPS complex size in the context of validated internal standards in (A) CA46 (left) and DG75 (right) cell lines and (B) P493-6 cells containing tetracycline (MYC OFF, left) or lacking tetracycline (MYC ON, right). (C) NADP+ levels, (D) [NADPH]/[NADP+] ratio, (E) NADH levels, (F) NAD+ levels, (G) [NADH]/[NAD+] ratio, (H) oxidized glutathione (GSSG) levels, (I) total glutathione levels and (J) [GSH]/[GSSG] ratio in WT, PRPS1- and PRPS2-KO cells of CA46 (left) and DG75 (right) cell lines, measured via relative luciferase units (RLU) of luminescent-based GLO-assays. in WT, PRPS1- and PRPS2- KO cells of CA46 (left) and DG75 (right) cell lines. For all panels, statistical analysis performed via One-Way ANOVA, bars represent mean ± s.d.; *p<0.05, **p<0.01, ***p<0.001, ****p<0.0001, ns: not significant.

**Supplementary Figure 3 – Related to Figure 3.**
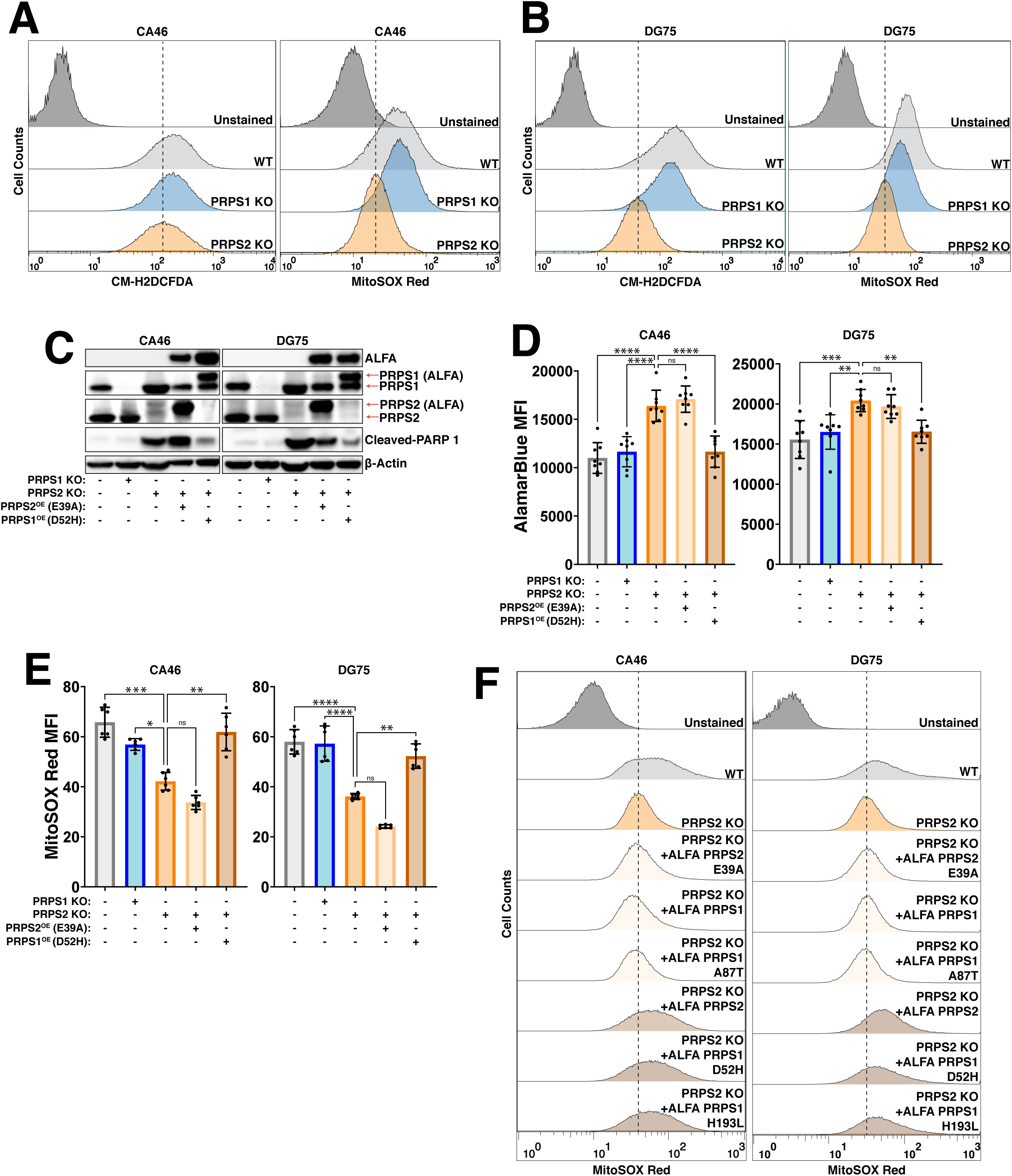
ROS accumulation in WT, PRPS1- and PRPS2- KO cells of (A) CA46 and (B) DG75 cell lines, measured via the total intracellular CM-H_2_DCFDA (left) and mitochondrial-specific MitoSOX Red (right) dyes. (C) Western blot validating stable exogenous expression of ALFA-tagged PRPS1 superactive (D52H) and PRPS2 catalytically inactive (E39A) mutant constructs in CA46 (left) and DG75 (right) PRPS2 KO cell lines. 24 kDa PARP-1 fragment is used as an apoptotic marker. β-Actin used as a loading control. (D) AlamarBlue mean fluorescence intensity (MFI) as a readout of intracellular reduction and (E) MitoSOX Red MFI as a readout of mitochondrial ROS accumulation in WT, PRPS1 KO, PRPS2 KO and PRPS2 KO cells containing stably integrated ALFA-tagged PRPS1 superactive mutant (D52H) and PRPS2 catalytically inactive mutant (E39A) constructs in CA46 (left) and DG75 (right) cell lines. (F) Mitochondrial ROS accumulation in WT, PRPS1 KO, PRPS2 KO and PRPS2 KO cells stably integrated with ALFA-tagged PRPS1, PRPS2, PRPS1 hypomorphic mutant (A87T), PRPS1 superactive mutant (D52H, H193L) and PRPS2 catalytically inactive mutant (E39A) constructs in CA46 (left) and DG75 (right) cell lines, measured via MitoSOX Red. For all panels, statistical analysis performed via One-Way ANOVA, bars represent mean ± s.d.; *p<0.05, **p<0.01, ***p<0.001, ****p<0.0001, ns: not significant. For all histograms, dashed lines indicate MFI of PRPS2 KO cells.

**Supplementary Figure 4 – Related to Figure 4.**
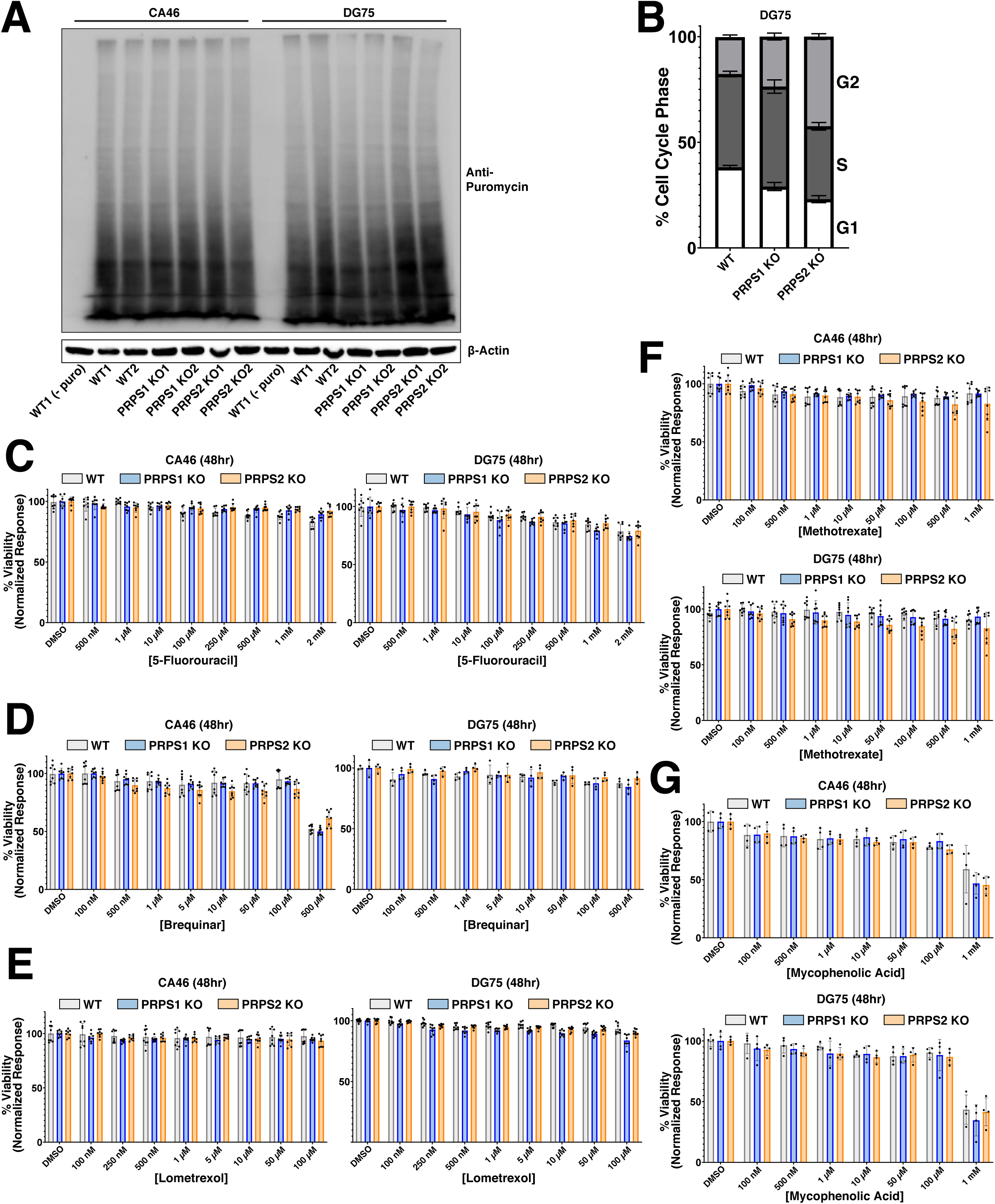
(A) Puromycylation assay measuring protein synthesis in WT, PRPS1- and PRPS2- KO cells of CA46 and DG75 cell lines. Each cell line is represented with two independent clones, per genotype. β-Actin used as a loading control. (B) Cell cycle analysis profiling the percentage of cells in G1, S, and G2 phases for WT, PRPS1- and PRPS2- KO cells of DG75 cells (C-G) Viability response of individual replicates of WT, PRPS1- and PRPS2- KO cells of CA46 (left/top) and DG75 (right/bottom) cell lines to treatment with increasing concentrations of (C) 5-fluorouracil, (D) brequinar, (E) lometrexol, (F) methotrexate and (G) mycophenolic acid, normalized to vehicle treatment. For all panels, bars represent mean ± s.d.

**Supplementary Figure 5 – Related to Figure 5.**
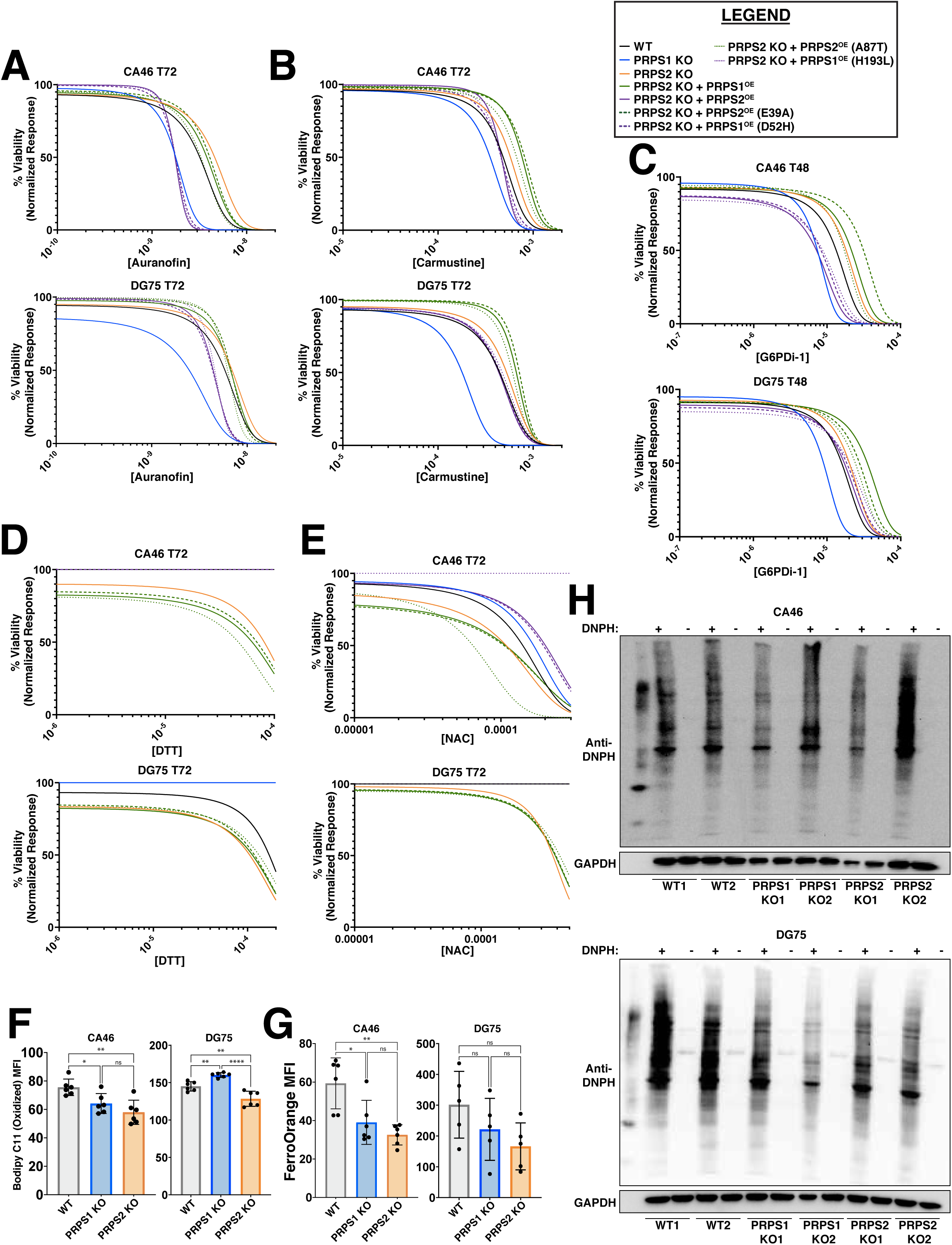
(A-E) Dose-response curves illustrating viability responses to (A) auranofin, (B) carmustine, (C) G6PDi-1, (D) DTT and (E) NAC treatment in WT, PRPS1 KO, PRPS2 KO and PRPS2 KO cells stably integrated with ALFA-tagged PRPS1, PRPS2, PRPS1 hypomorphic mutant (A87T), PRPS1 superactive mutants (D52H, H193L) and PRPS2 catalytically inactive mutant (E39A) constructs in CA46 (top) and DG75 (bottom) cell lines, normalized to vehicle treatment. X-axis represents the logarithmic scale of increasing drug concentration, Y-axis represents the normalized response as a viability percentage. Time points are indicated on each individual graph, determined by R^2^ values for goodness-of-fit (Data represented as a mean of normalized response of individual replicates at each concentration tested). (F) Lipid peroxidation, measured via BODIPY C11 mean fluorescence intensity (MFI) and (G) labile intracellular iron, measured via FerroOrange MFI in WT, PRPS1- and PRPS2- KO cells of CA46 (left) and DG75 (right) cell lines. (H) Western blot illustrating levels of global protein oxidation, via carbonyl side chain derivatization by 2,4-dinitrophenylhydrazine (DNPH), in WT, PRPS1- and PRPS2- KO cells of CA46 (top) and DG75 (bottom cell lines). (-) DNPH lanes serve as a control for DNPH-mediated derivatization. GAPDH is used as a loading control. For all panels, statistical analysis performed via One-Way ANOVA, bars represent mean ± s.d.; *p<0.05, **p<0.01, ****p<0.0001, ns: not significant.

